# Robust and accurate diagnosis of infectious skin diseases from histopathology images by integrating deep learning and explainable AI

**DOI:** 10.1101/2025.10.17.682660

**Authors:** Kpetchehoue Merveille Santi Zinsou, Habone Ahmed Mahamoud, Abdou Magib Gaye, Idy Diop, Maodo Ndiaye, Doudou Sow, Cheikh Talibouya Diop, Dmitry Korkin

## Abstract

Accurate diagnosis of infectious skin diseases remains a major challenge, particularly for neglected tropical diseases such as mycetoma, where precise pathogen identification is crucial for effective treatment. Histopathology imaging is the diagnostic gold standard, involving examination of tissue biopsies to identify characteristic inflammatory patterns, cellular changes, or microbial pathogens. However, its analysis is often limited by variability in tissue sampling and staining, subjective interpretation, inter-observer differences, and the absence of visible microbial grains in early disease stages. To elevate these challenges, we develop the Skin INfectious Diseases Intelligent (SINDI) framework, an integrated machine learning pipeline combining shallow learning, deep learning, stain normalization, and explainable AI to automate and enhance diagnostic accuracy from histopathology images. The SINDI framework is designed to systematically tackle increasingly complex tasks in diagnostics, including (1) disease phenotype classification and pathogen species identification, (2) understanding the importance of disease-specific regions (grains) and classification of grain-free images lacking visible microbial structures, (3) semantic segmentation of pathological features, and (4) explainable AI-driven interpretable decision support. Leveraging a comprehensive dataset of 1,324 histopathology images representing four predominant mycetoma pathogens that are curated by expert pathologists, alongside 7,000 healthy skin tissue images, SINDI demonstrated near-perfect accuracy in binary and multi-class classification tasks, particularly when employing Macenko stain normalization and domain-specific features. Remarkably, SINDI achieved high accuracy on images with masked grain regions and even on grain-free images, which are considered diagnostically intractable by human experts. Semantic segmentation models accurately delineated phenotype-related regions, while explainable AI methods provided transparent and clinically relevant interpretability of model decisions. Our results indicate that diagnostically relevant information is distributed beyond visible lesion areas, challenging traditional pathology paradigms. The SINDI framework thus represents a significant advance in automated infectious skin disease diagnostics, offering robust, interpretable, and scalable decision-support tools adaptable to diverse clinical settings.

## INTRODUCTION

Skin diseases present a significant global health burden [1], [2], affecting hundreds of millions of people worldwide with a wide spectrum of clinical severity, from minor self-limiting conditions to chronic, debilitating skin infections. While well-studied non-infectious skin diseases [3], such as psoriasis [4], [5], atopic dermatitis (eczema) [5], and melanoma [6], benefit from established diagnostic and treatment protocols, many infectious skin diseases disproportionately impact underserved communities with limited access to timely diagnosis and effective care. The latter diseases, many of which are categorized as Neglected Tropical Diseases (NTDs), highlight the critical need for accessible and automated diagnostic tools that can operate at the point of care, addressing the persistent shortage of specialized medical expertise in underserved regions [7], [8], [9], [10]. Developing such tools could greatly improve public health outcomes by enabling early and more accurate diagnosis at scale in resource-limited settings.

One of such examples of understudied but widely prevalent infectious skin diseases is mycetoma. It is a chronic and debilitating infectious disease that primarily affects subcutaneous tissues, caused by either fungal (*eumycetoma*) or bacterial (*actinomycetoma*) pathogens [11], [12]. Mycetoma is primarily contracted through traumatic skin injuries, and thus is prevalent in rural, socioeconomically disadvantaged communities, manual laborers, farmers, and those who walk barefoot, reflecting the disease’s strong association with poverty and occupational exposure. The disease is highly endemic within the “Mycetoma belt” [13], a band of countries spanning Africa, Latin America, and Asia. Characterized by painless subcutaneous masses, multiple fistulas, and the formation of visible grains containing the causative agent [14], mycetoma can lead to severe complications, extensive deformities, and functional loss if left untreated. Over 70 species of fungi and bacteria have been identified as etiological agents, which complicates both diagnosis and treatment [15], [16], [17]. Early diagnosis and treatment are crucial in reducing morbidity; however, the disease’s slow progression and indolent nature often lead patients to present at advanced stages, making intervention more challenging.

While various diagnostic techniques have been developed over time [18], [19], *histopathology* remains the most widely adopted diagnostic approach in resource-limited regions and countries, like Senegal and Sudan, due to its cost-effectiveness and accessibility [20]. However, histopathological diagnosis presents challenges. Histopathology imaging data on mycetoma available to the scientific community is sparse, with efforts to establish systematic data collection being recently made [21], [22]. Another key challenge in working with the histopathology slides is that they are highly susceptible to variability arising from differences in staining techniques (*e.g.*, Hematoxylin and Eosin, H&E [23], [24]), tissue preparation protocols [25], and the diverse morphological presentations of grains across various species and disease stages [26]. Furthermore, a major hurdle arises when mycetoma grains are absent in tissue samples, or when they are misidentified as host tissue reactions or processing artifacts [26]. In such ambiguous instances, pathologists must resort to supplementary, often more expensive and time-consuming techniques like grain culture or molecular methods, which introduce additional diagnostic complexity and delays. Beyond technical inconsistencies, histopathological diagnosis heavily relies on the expertise and subjective interpretation of individual pathologists. This variability in training and experience can lead to diagnostic inconsistencies, misdiagnosis, and delayed treatment, especially when analyzing complex mycetoma cases [26]. This challenge is particularly pronounced in regions with limited access to trained pathologists, further underscoring the need for more standardized and automated diagnostic tools that can improve diagnostic reproducibility and reduce human subjectivity.

The application of Artificial Intelligence (AI) and Machine Learning (ML) to histopathology images has become a transformative field, moving beyond traditional image analysis to provide automated, objective, and quantitative insights [27], [28]. The majority of these efforts have focused on cancer diagnosis and prognosis, a domain where large, well-annotated datasets are increasingly available [29], [30], [31], [32], [33]. Early approaches often relied on traditional, “shallow”, ML techniques, such as Support Vector Machines (SVMs), k-nearest neighbors (k-NN), and Random Forests (RF), which required manual feature extraction from images, including texture, color, and shape descriptors [34], [35], [36], [37]. These methods, while effective for specific, well-defined tasks, were limited by their reliance on the expert-designed features and often lacked the capacity to generalize across different datasets or diseases. Methods that employ image-related features, such as Haralick texture descriptors [38] or color histograms [39], have been used to classify breast cancer subtypes with moderate success (k-NN: 97.6%, SVM: 95.8%, and RF: 95.3% accuracy). However, higher accuracy rates of 98.25% with Random Forests and 93.5% with SVM were also achieved for breast cancer classification by segmenting tumor regions and extracting nine categories of features [40].

The paradigm shifted with the advent of deep learning, particularly Convolutional Neural Networks (CNNs) [41], [42], [43]. CNNs benefit from automatically learning hierarchical, discriminative vector representations extracted directly from raw image pixels, bypassing the need for manual feature engineering. Earlier work in this area has demonstrated strong performance on tumor detection, grading, and even inference of genetic variations from histopathology slides [44]. Models leveraging deep learning architectures such as ResNet, VGG, or EfficientNet, have routinely outperformed shallow learning methods [45], [46], [47], [48], [49], [50] and in several settings have achieved pathologist-level accuracy on specific benchmarks [45], [51], [52]. For example, one study [53] reported 97.8% accuracy for breast cancer using an autoencoder within a Siamese framework, while another [54] achieved 98.0% accuracy on skin cancer detection with an EfficientNetV2 model. Similarly, a Deep Residual Attention Network (DRANet) [55] was developed to classify 11 types of skin cancer, reaching an accuracy of 86.8% and providing visual rationales for its decisions. Despite these advancements, the limited interpretability (“black-box”) of these models continues to hinder clinical adoption, where transparency and reliability are essential. Moreover, as noted by others [56], [57], [58], the state-of-the-art systems demand large, well-annotated, and carefully processed datasets for training and are computationally expensive, complicating deployment in real-world settings.

In contrast to the extensive research on cancer histopathology, the application of machine learning to infectious skin diseases is a nascent but rapidly growing area. Several challenges persist: limited availability of infectious dermatopathology data, pronounced variability across pathogenic agents, and a lack of standardized imaging protocols in the field. In addition, most existing ones focus on a small number of well-characterized pathogens. For example, studies explored the use of CNN-based frameworks for detecting Leishmaniasis from Giemsa-stained microscopic slides using object detection methods [59], while others developed LeishFuNet by fine-tuning pre-trained CNNs to identify Leishmania parasites with limited image datasets [60]. Additional efforts include deep learning approaches to classify fungal species in microscopic images, reducing diagnostic delays and reliance on biochemical assays [61]. Nevertheless, there remains a substantial gap in developing a robust, generalizable framework capable of handling a wide range of infectious skin diseases while providing clinically actionable insights, particularly for neglected tropical diseases such as mycetoma.

Successful application of ML to histopathology is critically dependent on mitigating the technical variability inherent in slide preparation. Differences in staining protocols, reagent lots, and scanner settings can introduce significant color variations across different slides, even from the same lab, which can degrade the training and, as a result, predictive accuracy of a machine learning model [62]. To address this, advanced image normalization techniques have been developed and successfully used to standardize the appearance of histopathology images, making models more robust and generalizable [63]. While basic techniques, such as color space transformations or histogram matching [63], can correct global color shifts, more sophisticated methods are required to address the complex nature of histopathology staining. Three techniques in digital pathology have shown particular promise. Macenko stain normalization models the staining process as a linear combination of two stain vectors (*e.g*., Hematoxylin and Eosin) and a concentration matrix [64]. By estimating these components, it standardizes the image to a reference, removing variations in stain intensity and color without losing critical structural information [58], [65], [66], [67]. A more recent technique, Vahadane stain normalization, uses non-negative matrix factorization to decompose the image into stain colors and a corresponding concentration map [68]. This method offers a highly effective way to correct both stain intensity and color variations while preserving biological interpretability, improving model generalizability across diverse hospital datasets [69], [70]. Finally, the Reinhard stain normalization method is a robust and widely used statistical approach that transforms the color distribution of a source image to match a reference image [71]. This simple yet effective approach is computationally efficient and a popular choice for achieving consistent color appearances in computational pathology [72], [73].

Here, we address the pressing clinical and technical challenges of studying infectious skin diseases by introducing the Skin INfectious Disease Intelligent (SINDI) framework. The framework leverages state-of-the-art shallow learning and advanced deep learning architectures as well as novel explainable AI methods for addressing several key diagnostic tasks: determining a disease phenotype, identifying the causative pathogenic agents, and assisting the pathologists in the interpretation of the results. Here, we demonstrate the utility of the SINDI framework by applying it to study mycetoma, with a goal of developing a computational platform capable of not only accurately classifying mycetoma species but also providing transparent, clinically interpretable insights into their decision-making processes. We investigate if and for which diagnostic tasks an automated machine learning approach can match or even surpass human expert capabilities. We also study the role of stain normalization in unlocking subtle diagnostic information and determine if AI can leverage features beyond human visual perception. Ultimately, our work aims to reduce the diagnostic burden on pathologists, particularly in regions with limited access to expertise, and to provide clinicians with a powerful, trustworthy, and efficient tool for rapid, accurate, and species-specific diagnosis of infectious skin disease and treatment planning.

## RESULTS

In this work, we developed and systematically evaluated the Skin INfectious Disease Intelligent (SINDI) machine learning-based framework across five core tasks aimed at addressing persistent challenges in the diagnosis of infection skin diseases: (1) classification of disease phenotype and pathogenic species causing the infection, (2) analysis of the impact of grain-like regions indicative of phenotype (Regions of Interest, ROIs) on classification accuracy, (3) classification of disease phenotype and pathogenic species using grain-free histopathology images (*i.e.*, images without identifiable ROIs), (4) semantic segmentation of histopathology images into pathologically salient substructures, and (5) explainable AI-guided interpretation of the classification results by a ML model. The tasks are designed to be increasingly difficult: from standard diagnostic tasks (Task 1), comparable to expert pathologist practice, to highly challenging scenarios (Task 3), which are nearly impossible for human experts due to the absence of diagnostically relevant features, and to Task 5, where the framework guides an expert through a decision-making process by a machine learning classifier. The SINDI framework was assessed on a large-scale curated dataset of 1,324 mycetoma histopathology images, which included cases caused by two bacterial and two fungal pathogens. Together, these tasks form an integrated diagnostic pipeline that combines image preprocessing, shallow and deep machine learning, and explainable AI-guided interpretability techniques. Across the spectrum, from conventional species classification to the more complex challenge of grain-free image analysis, our framework consistently achieved performance that matched or surpassed that of expert pathologists.

### A comprehensive multi-pathogen histopathology data of skin disease

The dataset collected for the supervised learning and explainable AI tasks is an expanded dataset of 1,324 histopathology images representing four predominant mycetoma-causing pathogens: *Actinomadura pelletieri (AP)*, *Falciformispora senegalensis (FS)*, *Madurella mycetomatis (MM)*, and *Streptomyces somaliensis (SS)* (Fig. 1, 2a). First, these images were labeled by two expert pathologists to categorize them into their mycetoma species. Next, the same two expert pathologists independently annotated each image to generate two distinct sets of ‘ground truth’ masks (Mask1 and Mask2, Fig. 1), emphasizing the grain regions and ensuring robustness against inter-observer variability. A mask represents a union of all isolated parts of an image, each part corresponding to a grain region. In total, 33 distinct image datasets were generated for the five tasks described below and used for model training, evaluation, and analysis across these tasks (Supp. Fig. S1, Supp. Fig. S2).

**Figure 1.**
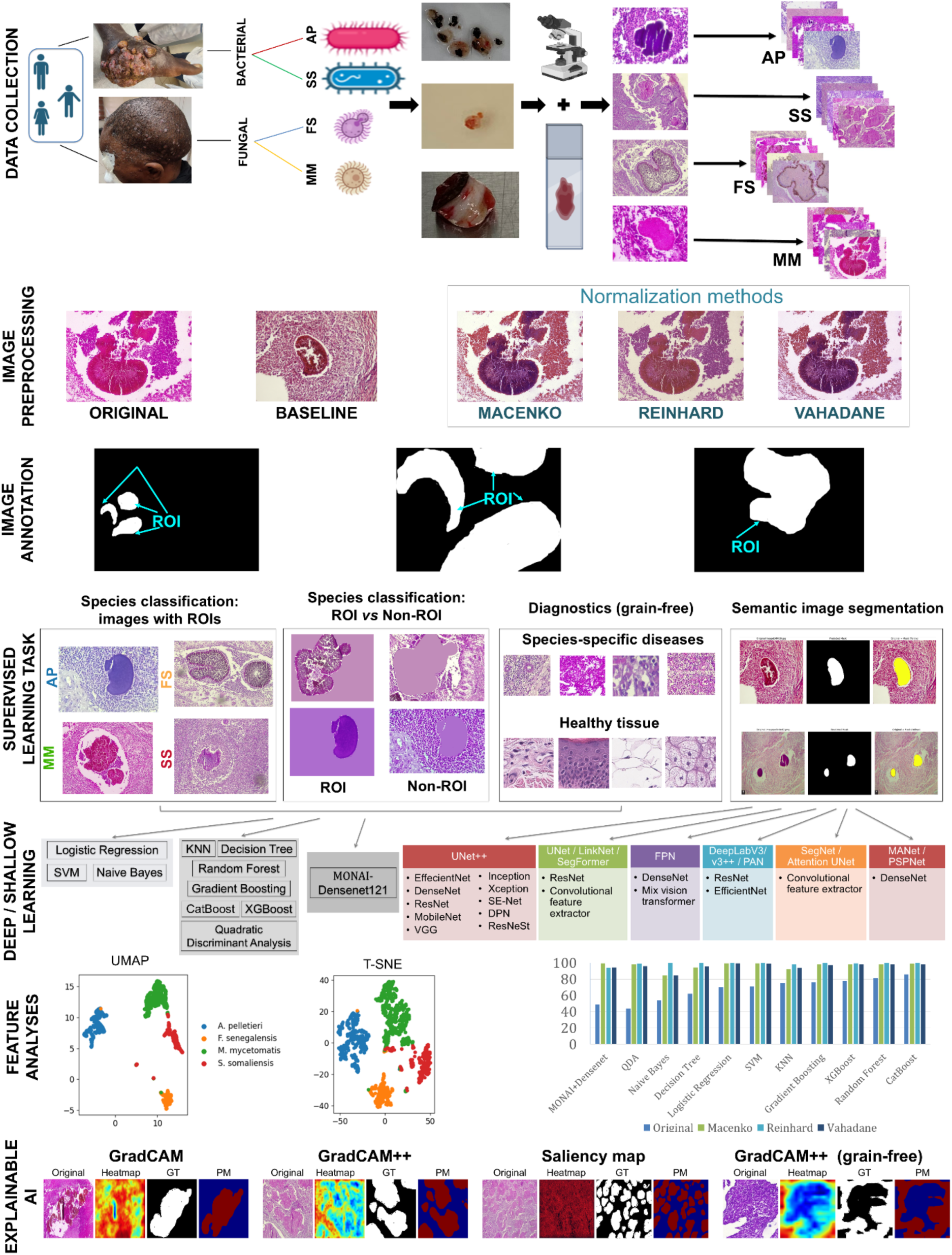
Overview of the SINDI Framework for Mycetoma Histopathology Analysis. **1. Data Collection**. Stained histopathology images of mycetoma samples, representing four prevalent bacterial and fungal species from Senegal, were acquired using a Leica microscope, followed by expert-based annotation of the pathogenic agents causing the phenotype. **2. Image Preprocessing.** Original images were normalized to a baseline using Macenko, Reinhard, and Vahadane methods to improve their consistency. **3. Image Annotation.** Grain-rich Regions of Interest (ROIs, colored white) annotations were generated using Label Studio, producing ground-truth masks. **4. Supervised Learning Tasks.** Shown are representative examples of the four distinct supervised learning tasks performed within the framework: (i) Standard disease phenotype and species classification using histopathology images enriched with ROIs; (ii) Analysis of the role of ROIs in the classification where the histopathology images have ROIs masked or only ROIs present; (iii) disease phenotype and species classification using grain-free histopathology images, *i.e.,* without any visible ROIs; and (iv) semantic image segmentation into histopathologically salient regions. **5. Deep/Shallow Learning Models Evaluation and Optimization.** Classification models are grouped into linear, non-linear, and deep learning categories. Deep learning models for semantic segmentation are listed with their various encoder architectures. **6. Feature Analysis:** UMAP and t-SNE embedding showcase learned feature representations. A comparative analysis of classifications using raw images versus the images normalized with one of the three normalization techniques is carried out. **6. Explainable AI:** Results from Grad-CAM, Grad-CAM++, and Saliency-based gradient explainable AI methods are displayed, showing the original image, saliency heatmap (continuous values), manual ground-truth (GT), and binary predicted mask (PM) for each method.

For Task 1, consisting of two subtasks, we generated eight distinct datasets (Supp. Fig. S1a). For subtask 1 (binary classification of disease and healthy tissues), we evaluated the 1,324 disease images and 7,000 healthy tissue images from the Skin Tumor Histology Dataset [54]. For subtask 2 (multi-class classification of four pathogenic species), the dataset of 1,324 disease images was used. Both subtasks included classifications based on raw, Macenko-normalized, Reinhard-normalized, and Vahadane-normalized images. For Task 2 (ROI *vs*. Non-ROI Impact on Species Classification), we created eight paired datasets (Supp. Fig. S1b). These datasets consisted of 1,324 ROI-isolated images and 1,324 Non-ROI images (see Methods for more details), generated from both original and Macenko-normalized images, and using both Mask1 and Mask2 annotations. For Task 3 (Grain-Free Image Classification), we generated eight distinct datasets for evaluation (Supp. Fig. S1c). Specifically, we included 3,034 or 3.027 grain-free image fragments derived from the original specimens (using Mask1 and Mask2 annotations, respectively, for both raw and Macenko-normalized versions). Models were evaluated on these fragments both independently and in combination with the 7,000 healthy tissue images as negative controls, forming the eight datasets. This comprehensive dataset design enabled rigorous evaluation across various diagnostic scenarios, from clear ROI cases to challenging grain-free specimens, while maintaining robust controls to assess model specificity and the spatial distribution of diagnostic features. For Task 4 (Semantic Image Segmentation), the manual ground truth annotations (Mask1 and Mask2) served as the primary datasets, resulting in two distinct ground truth sets for model training and evaluation (Supp. Fig. S2a). For Task 5 (Explainable AI for Diagnostic Decision Support), seven datasets of heatmaps were generated (Supp. Fig. S2b). Six datasets involved combinations of raw and Macenko-normalized images, derived using Grad-CAM, Grad-CAM++, and Saliency gradient-based techniques. The seventh dataset contains the generated heatmap for grain-free images using Grad-CAM++.

### Evaluation of Task 1 showed that disease-causing pathogen identification is a more challenging problem than a disease *vs.* healthy classification

Task 1 was the easiest to solve, essentially replicating diagnostic tasks performed by the pathologist (Fig. 1, Fig. 2a). In this task, we also aimed to evaluate the effectiveness of various feature extraction strategies and stain normalization techniques. In the first subtask, aiming to distinguish between healthy tissue and mycetoma-infected samples, we compared domain-independent and domain-specific feature-based models, as well as a deep learning model strategy (Fig. 2b, d, Supp. Fig. S3a, b). Given a substantially large dataset of labeled examples (1,324 disease images alongside 7,000 healthy histopathology images), one would expect to get a good performance from both shallow learning and deep learning models. We also tested the impact of three different preprocessing methods, Macenko, Reinhard, and Vahadane. These image preprocessing methods have been successfully introduced in the field of histopathology [64], [68], [71], but were never used to study infectious skin diseases; thus, it was not clear if these methods would improve the classification accuracy.

**Figure 2.**
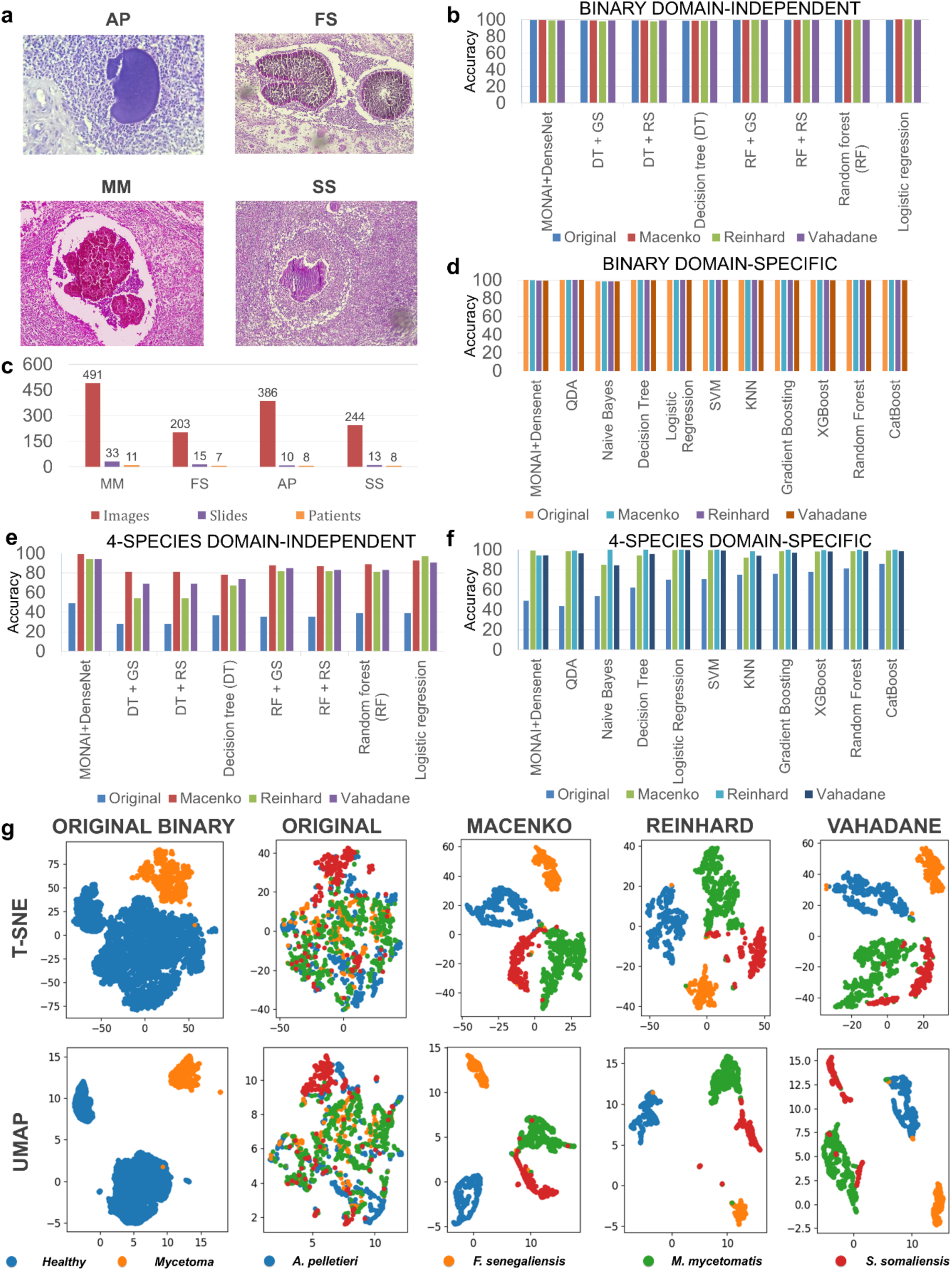
Analysis of model performances for Task 1 concerned with the classification of disease samples and the source pathogens. **(a)** Examples of histopathology images for each of the four mycetoma species analyzed. **(b)** Binary disease classification performance using unprocessed and normalized image datasets and domain-independent features. Training results of four classifiers, including three shallow learning classifiers with varying optimization parameters (Decision Tree (DT), Random Forest (RF), and Logistic Regression), and one deep learning (MONAI+DenseNet) model. The methods are ordered based on their performance on the original image dataset. **(c)** Dataset Composition. Detailed distribution of collected images, slides, and patients across four species. **(d)** Binary disease classification performance using unprocessed and normalized image datasets and domain-specific features. Training results of 11 classifiers, including ten shallow learning and one deep learning (MONAI+DenseNet) models. The methods are ordered as in (b). **(e)** Multiclass species classification performance using unprocessed and normalized image datasets and domain-independent features. Training results of the same four classifiers defined in (b). The methods are ordered as in (b). **(f)** Multiclass species classification performance using unprocessed and normalized image datasets and domain-specific features. Training results of 11 classifiers, including ten shallow learning and one deep learning (MONAI+DenseNet) models. The methods are ordered as in (b). **(g)** t-SNE and UMAP based feature embeddings for two-class (diseases vs. healthy) and four-class (pathogenic species) groupings of the original and normalized image datasets. In the four-class groupings, the embedding shows a substantially better separation of the data points corresponding to images processed with either of the three normalization methods.

For the first subtask, using domain-independent approaches, Random Forest achieved 99.69% accuracy on the raw images, improving to 99.87% with Macenko normalization, while Reinhard and Vahadane showed comparable performance at 99.39% and 99.87%, respectively (Fig. 2b). The domain-specific approach for the same subtask proved to be the most robust, with 9 out of 10 implemented and trained supervised classifiers achieving perfect or near-perfect classification (100%) across all three preprocessing methods and even on the raw data (original images). Interestingly, the deep learning model (MONAI + DenseNet121) performed similarly, achieving up to 99.94% accuracy on the original images, though slight variations were observed across stain normalization methods (99.76% Macenko, 99.76% Reinhard, 99.28% Vahadane) (Fig. 2b).

The second subtask focused on a more challenging problem of classifying a pathogen causing the infection. Here, the goal was to classify mycetoma cases by labeling them into one of four causative pathogenic species using the same training/testing set of 1,324 labeled images. As with the previous task, we evaluated domain-independent, domain-specific, and deep learning methods (Fig. 2e,f, Supp. Fig. S3a,b, Supp. Fig. S7). As a result, we observed that the domain-independent models using raw images performed poorly (*e.g*., Random Forest at 39% accuracy, Fig. 2e), but accuracy significantly improved with stain normalization, reaching up to 89%, 81%, 83% when using Macenko, Reinhard, and Vahadane preprocessing methods, respectively (Fig. 2e). Deep learning showed an even greater sensitivity to preprocessing: MONAI+Densenet121 achieved only 49% on raw images, but spiked to 99.34% with Macenko-normalized images (Fig. 2e,f). Domain-specific approach consistently outperformed the others in this subtask, with several classifiers (CatBoost, Random Forest, XGBoost) achieving near-perfect results (up to 100%) using Reinhard and Macenko-normalized images (Fig. 2f). Overall, the evaluation of the classification performances in subtask 2 confirmed that for species classification, domain-specific features, and stain normalization are more critical to achieve high accuracy than for subtask 1.

To visualize the trends observed during the classifications, we applied two commonly used feature embedding approaches, UMAP and t-SNE, to the vector representations of the labeled examples using domain-specific features (Fig. 2g, Supp. Fig. S4, Supp. Fig. S6). In the disease classification subtask, the visualization revealed a strong grouping of the healthy and diseased samples, even when using raw images. This visual separation is in concordance with the high performance observed across all modeling strategies in binary disease classification. However, for multi-class species classification in Subtask 2, no clear groups were visible using the raw image features (Fig. 2g). After applying three different stain normalizations, a clearer species-level grouping appeared in the embedding maps (Fig. 2g).

### Evaluation of Task 2 showed that image preprocessing enables accurate classification independent of regions of interest

During the traditional diagnosis of infectious skin diseases using histopathology slides, pathologists primarily focus on the regions of interest (ROIs), grain-rich areas that are indicative of the disease phenotype. However, the visibility and diagnostic quality of these ROIs can vary considerably across images, leading to potential misdiagnoses or misclassification of pathogenic species [18], [26]. Task 2 (Fig. 1, Fig. 3a) was therefore designed to assess whether species classification accuracy depends strictly on these diagnostically salient regions, or whether sufficient information can also be extracted from the images, which had all ROIs removed. Specifically, we tested the performance of the classifier on the four-class pathogen classification problem (see subtask 2 in Task 1) using two types of images: one where everything but ROIs is ‘scrambled’ by replacing the original pixels in Non-ROI regions with the image-averaged values, and another one where all ROIs were similarly scrambled using the image-averaged values (Supp. Fig. S8a). We refer to the first group of images as ROI images and the second one as Non-ROI images. Lastly, we tested if the image preprocessing methods could improve the classification accuracy for either type of the images.

**Figure 3.**
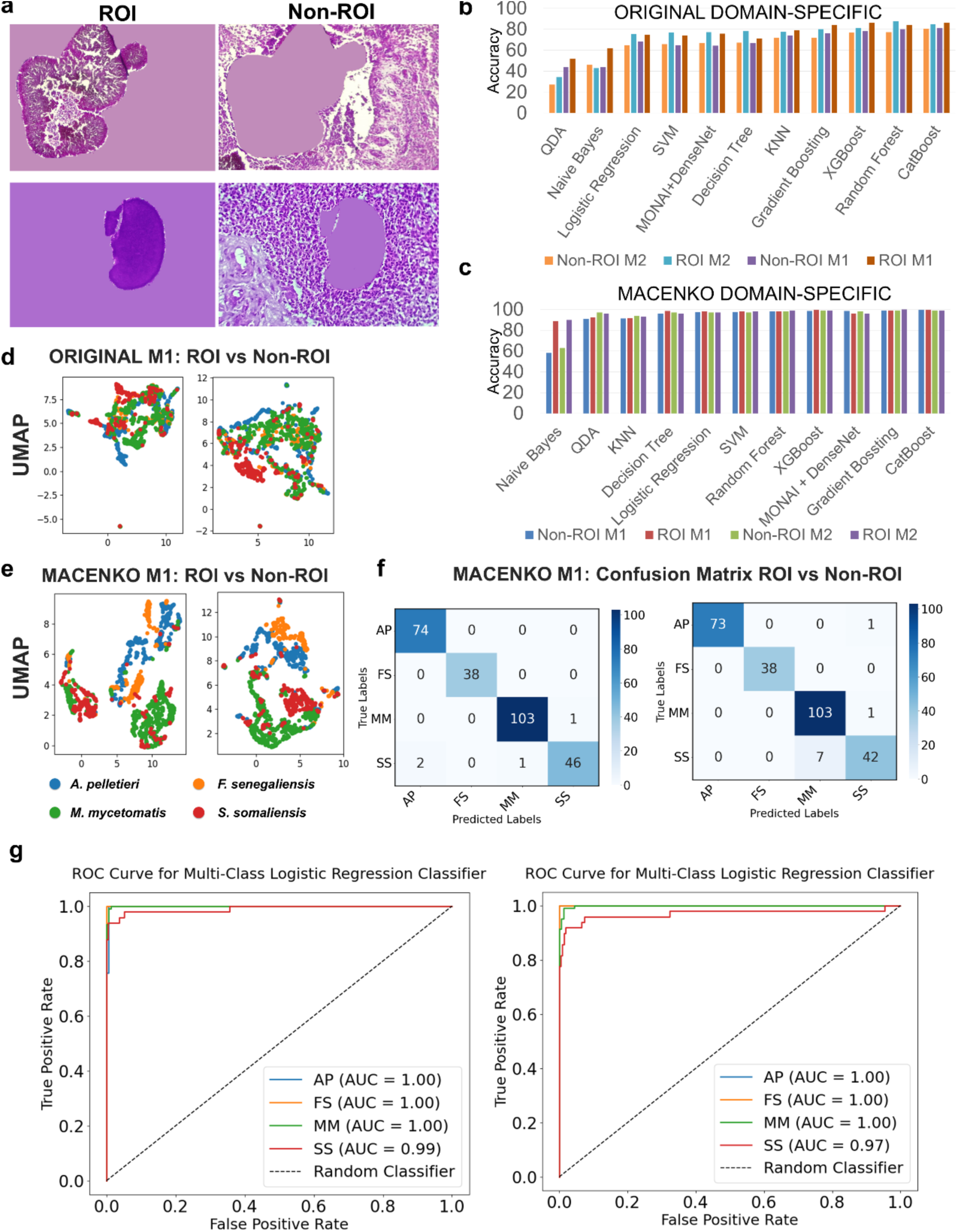
Analysis of model performances for Task 2 concerned with the assessment of the impact of ROIs and Non-ROIs present in the images on species classification accuracy. **(a)** Examples of ROI and Non-ROI images. The ROI images have only ROIs visible, while everything else is masked using an averaged color; Non-ROI images have ROIs masked. **(b)** Classification performance using unprocessed images with expert-annotated ground truth masks, Mask1 and Mask2, curated by two expert pathologists, respectively. Evaluation results of ten classifiers comparing domain-specific feature representation for shallow learning *vs.* a deep learning (MONAI+ Densenet) approach. The methods are ordered based on their performance for the most difficult Non-ROI dataset using Mask1. **(c)** Classification performance using images normalized with Macenko method and with expert-annotated ground truth masks, Mask1 and Mask2, curated as in (b). Evaluation results of ten classifiers comparing domain-specific feature representation for shallow learning *vs.* a deep learning (MONAI+ Densenet) approach. The methods are ordered using the same criterion as in (b). **(d)** UMAP-based feature embedding for ROI versus Non-ROI datasets generated from original, unprocessed images using Mask1 (M1). **(e)** UMAP-based feature embedding for ROI versus Non-ROI datasets generated from images processed with Macenko normalization method using Mask1 (M1). The embedding shows a substantially better separation of the data points corresponding to the individual pathogens, compared to (d). **(f)** Multiclass species classification performance for ROI versus Non-ROI datasets, based on Macenko-normalized images and Mask1 characterized by confusion matrices. **(g)** One-against-all species classification performances for a multi-class logistic regression classifier, for ROI and Non-ROI datasets, based on Macenko-normalized images and Mask1, characterized by the Receiver Operating Characteristic (ROC) curve.

Building on assessment results from Task 1, which showed consistent underperformance by domain-independent methods, we focused exclusively on the remaining two strategies: domain-specific feature extraction and the MONAI-based DenseNet deep learning framework. The ROI and Non-ROI areas were then extracted from the original and Macenko-normalized datasets. The Macenko preprocessing method was again selected due to its superior performance over two other preprocessing techniques in Task 1. The ROIs and Non-ROIs were generated using manually annotated ground truth masks, Mask1 (M1) and Mask2 (M2), from the two pathologists, resulting in four distinct datasets: ROI-M1, Non-ROI-M1, ROI-M2, and Non-ROI-M2, each containing the four mycetoma species.

When assessing the performances of multi-class classifiers on the original, unprocessed images, we found that for both annotation sets (M1 and M2), ROI-based models outperformed their Non-ROI counterparts (Fig. 3b, Supp. Fig. S8b), highlighting the importance of the diagnostically salient areas within the image. This observation is intuitively expected because grain-rich areas (ROIs) have long been used by pathologists as the most informative parts of the slide when forming a diagnosis. Overall, the performance varied substantially across the models and region types. Domain-specific approach with classifiers such as Random Forest, XGBoost, and CatBoost consistently outperformed deep learning models. CatBoost reached 86% accuracy on ROI-M1 and 84.5% on ROI-M2, while MONAI+DenseNet lagged slightly behind with accuracies of 75.75% (ROI-M1) and 77.15% (ROI-M2).

Unexpectedly, when the same experiment was conducted on Macenko-normalized image types, the distinction between ROI and Non-ROI based performances essentially disappeared (Fig. 3c,f,g, Supp. Fig. S8c), highlighting, for the first time, the superior diagnostic power of the classification methods over pathologist-based diagnostics. Models such as CatBoost, Gradient Boosting, and SVM achieved nearly identical and near-perfect accuracies across all region types (Fig. 3c, Supp. Fig. S8c). Gradient Boosting achieved 99–100% accuracy across ROI and Non-ROI, for both M1 and M2 annotations. MONAI+DenseNet also performed consistently in the high 95–98% range, though still slightly lower than the top-performing conventional models. We also observed that the performance over the four pathogens differed, although not significantly (Fig. 3f,g), potentially due to the small but important phenotypic differences. To visualize the distribution of learned features, we embedded the domain-specific feature vector space using UMAP and t-SNE. In both the original and Macenko datasets, the species-based grouping of data points was not clearly discernible, either for ROI or Non-ROI images (Fig. 3d,e, Supp. Fig. S8d).

### Successful ML-driven diagnostics based on grain-free images in Task 3, which are virtually unusable for expert-based diagnostics

In our final classification task (Task 3), we explored the limits of machine learning for pathogen-specific diagnostics in the most challenging clinical setting: grain-free histopathology images. The images did not have the visual evidence of fungal or bacterial colonies, and therefore no ROIs could be detected. Such cases are exceptionally difficult, often impossible, for a pathologist to resolve without complementary molecular or microbiological tests [74], [75]. Our aim was to determine whether a computational method, trained solely on grain-free images, could still extract discriminative features to differentiate between mycetoma species and between diseased versus healthy tissues.

To this end, we generated two types of datasets. First, we created grain-free datasets by dividing the original and Macenko-normalized images into fragments and discarding any that overlapped with annotated ROIs. Second, we extended the original and Macenko-normalized datasets by integrating 7,000 images of healthy skin tissue types from the Skin Tumor Histology Dataset. The types included dermis, epidermis, vessels, nerves, skeletal muscle, sebaceous glands, and subcutis (Fig. 1, Fig. 4a). These datasets were again constructed using two independent ground-truth annotation sets (Mask1 and Mask2) to ensure robustness (Supp. Fig. S9a).

**Figure 4.**
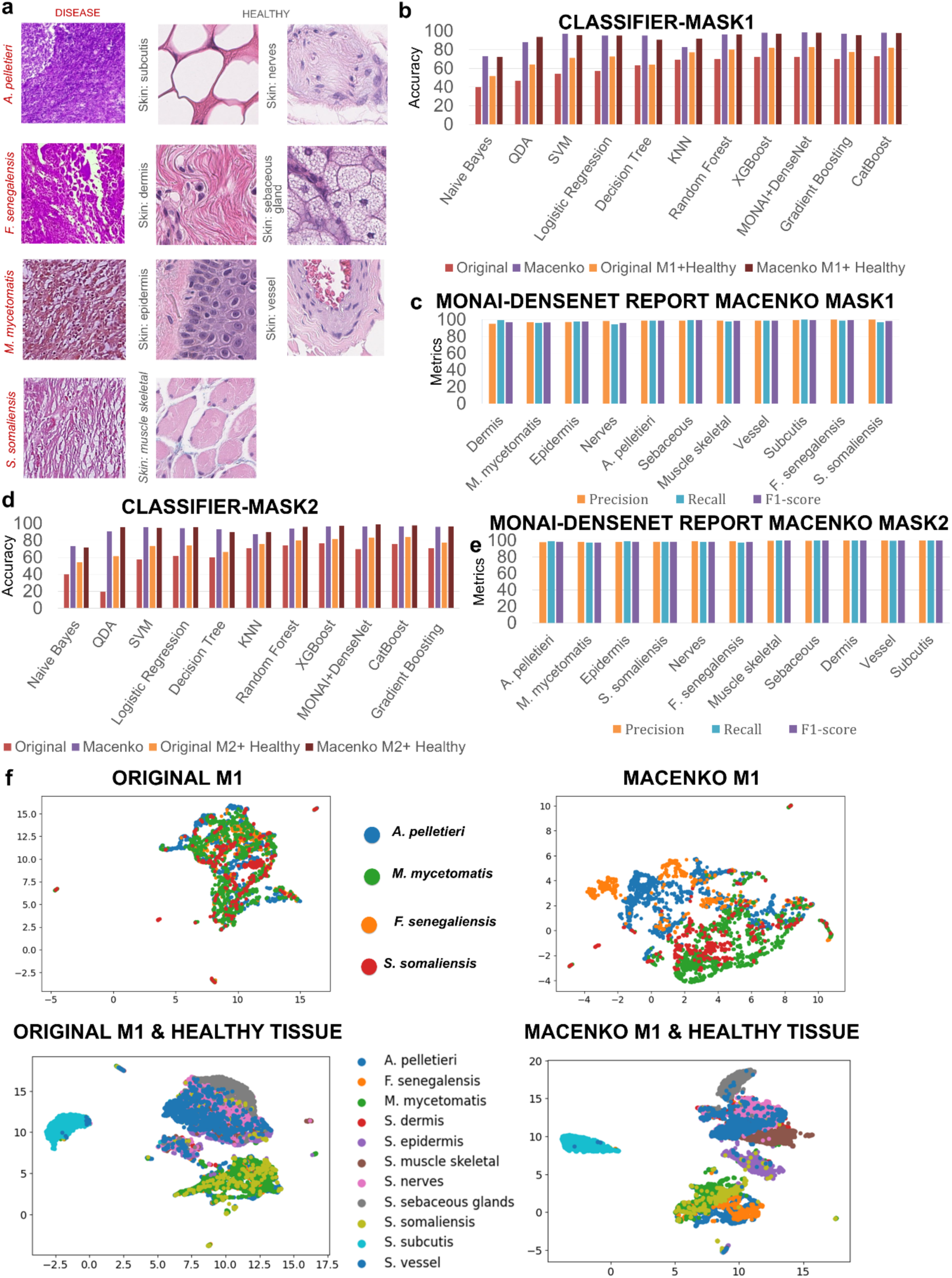
Analysis of model performances for Task 3 concerned with multiclass species classification for images with no visible grains. **(a)** Examples of grain-free histopathology images for each of the four mycetoma species analyzed and for seven healthy skin tissue types**. (b)** Multiclass classification performance for grain-free image datasets, based on Macenko-normalized images and a grain-free region generated based on Mask1 annotations. Evaluation results of ten classifiers comparing domain-specific feature representation for shallow learning *vs.* a deep learning (MONAI+ Densenet) approach on original grain-free, Macenko grain-free, and combined Macenko grain-free and healthy tissue datasets. The methods are ordered based on the accuracy values for the original dataset. **(c)** One-against-all classification performances on the pathogenic species and healthy tissue classes for MONAI+DenseNet model, using Mask1 annotations and Macenko normalization protocol. **(d)** Multiclass classification performance for grain-free image datasets as in (b) but using Mask2 annotations. **(e)** One-against-all classification performances on the pathogenic species and healthy tissue classes as in (c), but using Mask2 annotations. **(f)** UMAP-based feature embedding for original (top left) and Macenko-normalized (top right) grain-free datasets for the four pathogenic species, and original (bottom left) and Macenko-normalized (bottom right) datasets of pathogenic and healthy tissue classes. None of the embeddings was able to clearly partition the data into the corresponding classes.

We benchmarked species classification performance on grain-free images generated with each ground-truth mask by comparing 11 shallow machine learning classifiers using domain-specific features and a deep learning classifier (Fig. 4b-e, Supp. Fig. S9b, c). Each model was separately trained and evaluated on eight datasets: original grain-free images, Macenko-normalized grain-free images, and a combined dataset that integrated original images with healthy tissue images, Macenko-normalized images with healthy tissue images (four datasets in total for each, Mask1 and Mask2).

Our evaluation showed that the use of raw grain-free histopathology images is impractical for the classification tasks to identify individual pathogenic species. For the annotated dataset using Mask1, the most accurate classifier, MONAI+DenseNet, achieved only 72.5% accuracy, while closely followed by a shallow learning classifier, XGBoost, at 72% accuracy; for the dataset annotated using Mask2, XGBoost was the best performer at 76.6%. When classifying the four pathogenic species along with the seven skin tissues, the accuracy improved: the best accuracies for Mask1 and Mask2 annotations were 82.6% for MONAI+DenseNet and 83.8% for CatBoost, another shallow learning classifier, respectively (Fig. 4b,d, Supp. Fig. S9b). This is not surprising, because it is expected that classification between the disease and healthy skin tissues is easier than across the pathogen-specific disease tissues only.

Surprisingly, preprocessing the raw grain-free images using the Macenko method drastically improved the classification accuracy for both tasks and both annotations (Fig. 4b-e, Supp. Fig. S9b, c). The top-performing method for the four-pathogen classification task was MONAI+DenseNet (98.5% and 96.7% for Mask1 and Mask2 annotations, respectively), with XGBoost and CatBoost shallow classifiers trailing very closely. Similarly, the multi-class classification performance on the four pathogen-infected skin tissue types combined with the seven healthy skin tissue types achieved 98.2% and 99.1% accuracy by MONAI+DenseNet on Mask1 and Mask2 annotations, respectively (Fig. 4b-e, Supp. Fig. S9b, c). This again shows the critical importance of applying the correct preprocessing protocol to the raw histopathology images prior to machine learning tasks.

Finally, the visualization of the feature vectors generated for the raw and processed images using UMAP and t-SNE methods was also consistent with our understanding that this classification task is the most challenging one and that preprocessing helped to improve the separability of the data points from different classes (Fig. 4f, Supp. Fig. S10). Specifically, when applied to the raw grain-free images of four pathogenic classes, both embeddings showed no clear grouping. Furthermore, even after applying the Macenko normalization, only minor improvements in class separation were observed. The same behavior was observed when applying embeddings to the raw images of four pathogenic and seven healthy skin tissue types, with data points for mycetoma species and healthy tissue types intermixed. However, embedding of the Macenko-normalized images for the combined four pathogenic and seven healthy tissue classes showed drastic improvement in groupings based on the tissue type.

### Image segmentation and explainable AI methods in Tasks 4 and 5 provide new means in improving diagnostic quality and confidence by pathologists

Once our methods achieved outstanding prediction accuracy on a range of histopathology images and across different pathogens, an important question arose: How to “translate” the classification results so that they can be interpreted by pathologists more intuitively? Specifically, one would like to inform the pathologists about the reasons the model’s decision is made, so they can decide to trust the results or request a re-evaluation. To address this question, we implemented two different approaches: (1) a semantic image segmentation, where we structured each image by identifying its critical and phenotypically distinctive parts (Task 4), and (2) an explainable AI, where we introduced the ranking on the importance of the image’s regions towards the accurate classification by a ML classifier (Task 5). The two approaches are distinct in the way they deliver the information: while in Task 4, the method essentially identifies the regions of interest that characterize the phenotypic changes in the tissues, in Task 5, only those regions that are the most informative for the classifier are highlighted (which could be both regions of interest and other regions). We referred to these regions as saliency heat masks.

While the originally generated saliency heatmaps are continuous-valued, to further simplify the process of interpretation, these continuous-valued maps were converted to the binary saliency classification masks, specifying the regions of images that are important or non-important for the classification task.

For Task 4, the results of the automated segmentation using 12 different deep learning architectures with 13 pretrained encoder models (see Fig. 1, Fig. 5a-e, Supp. Fig. S11a, Supp. Fig. S12-14, see subsection ***Stand-alone deep learning models for semantic image segmentation task (Task 4)*** in Methods) were assessed on the two expert-annotated sets of images (Mask1 and Mask2). For the raw, unprocessed images, the classical deep learning architectures SegNet, UNet, and Attention UNet did not perform well on the Mask1 dataset, with Attention UNet reaching the Dice coefficient value of 86.9% and Intersection over Union (IoU) value of 77.0%, while UNet (Dice = 57.2%, IoU = 40.8%) and SegNet (76.0%, 62.0%) performed significantly worse (Supp. Fig. S12a). The introduction of the Macenko preprocessing stage led to substantial improvements for UNet (83.6%, 72.1%) and moderate gains for SegNet (81.3%, 69.0%). However, Attention UNet did not benefit from normalization, in fact experiencing a slight decline (86.2%, 76.0%) (Supp. Fig. S12a). Thus, these three architectures were not further evaluated on the Mask2 dataset.

**Figure 5.**
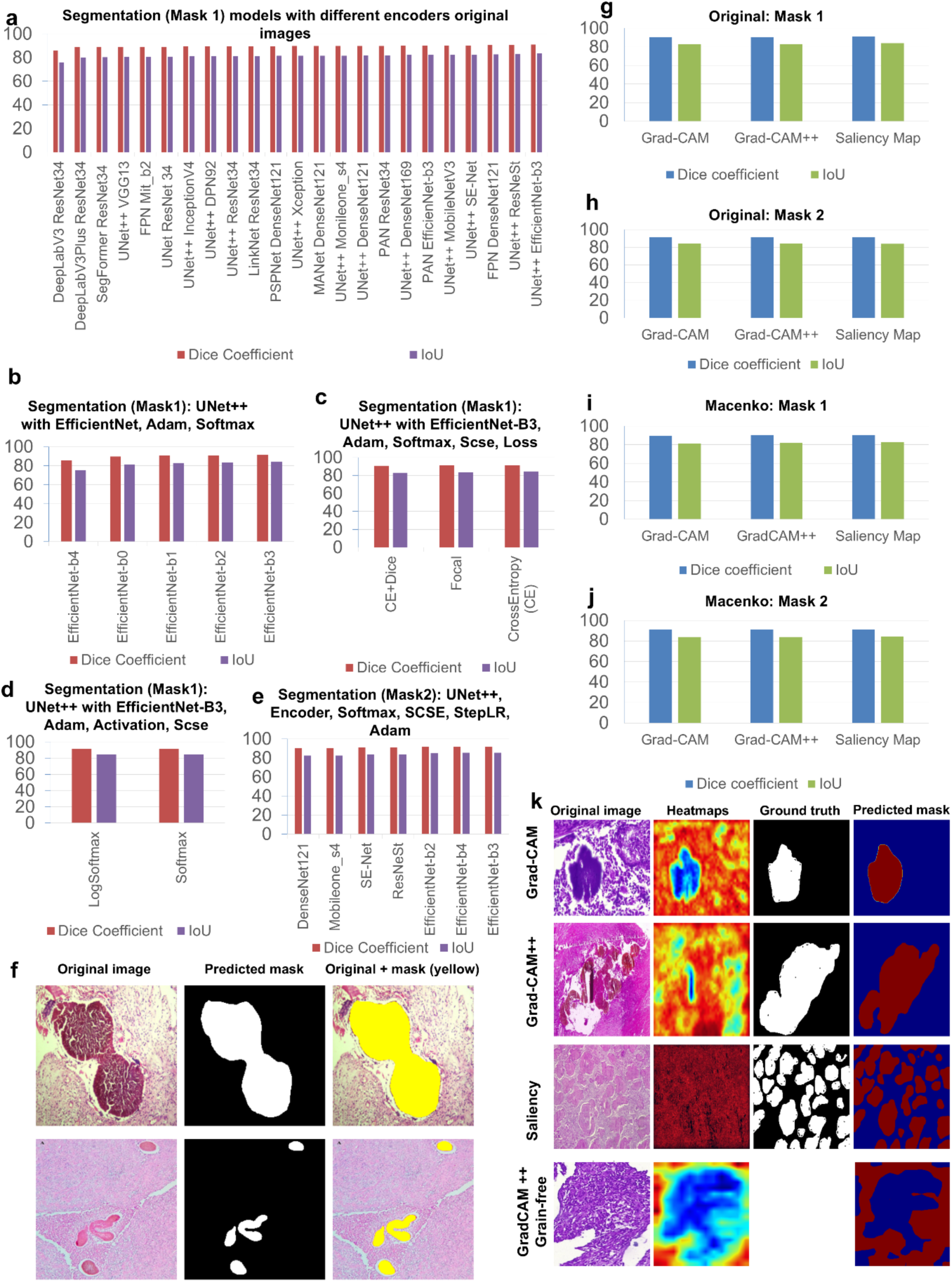
Analysis of model performances for Task 4 (Semantic Segmentation) and Task 5 (Explainable AI for Mycetoma Histopathology). **(a)** Performance assessments of 24 combinations of deep learning architecture and encoder models on Task 4, where the ground truth is defined using Mask1 annotations applied to the original images. For each of the 24 combinations, two metrics were calculated: Dice coefficient and IoU. **(b)** The performance of the deep learning model for semantic segmentation (Task 4) depends on specific architecture topology of the encoder. Performance evaluation of the top-performing architecture (UNet++ in combination with EfficientNet encoder, Adam optimizer, and Softmax activation) trained on original images with Mask1 annotations for each of the five architecture topologies: efficientnet-b0 – efficientnet-b4. **(c)** The performance of deep learning model for semantic segmentation (Task 4) depends on a specific loss function. Performance evaluation of the top-performing architecture (UNet++ in combination with EfficientNet encoder (efficientnet-b3 topology), Adam optimizer, and Softmax activation) trained on original images with Mask1 annotations for each of the three loss functions: CrossEntropy (CE), Focal, and CE combined with Dice. **(d)** The performance of deep learning model for semantic segmentation (Task 4) depends on a specific activation function. Performance evaluation of the top-performing architecture (UNet++ in combination with EfficientNet encoder (efficientnet-b3 topology), Adam optimizer, and CE loss function) trained on original images with Mask1 annotations for each of the two activation functions, softmax and logsoftmax. **(e)** The performance of deep learning model for semantic segmentation (Task 4) trained on original images with Mask2 annotations is comparable across different encoder models. **(f)** Example of a semantic segmentation model (Task 4) applied to a raw image. Shown are representative original images, their corresponding predicted segmentation masks, and overlays of the original images with predicted masks. **(g)-(j)** Evaluation of performances of three explainable AI models **(Task 5)** on original and Macenko-normalized datasets annotated with Mask1 and Mask2. **(k)** Example of three explainable AI models (Task 5) applied to a raw image with visible ROIs (first three rows) and the best-performing Grad-CAM++ model applied to a grain-free image (last row). Detailed examples of XAI outputs for each method (Grad-CAM, Grad-CAM++, Saliency), showing the original image, saliency heatmap (continuous values), manual ground-truth mask, and predicted saliency heat mask (binary values). An additional Grad-CAM++ example for a grain-free image is also included.

On the other hand, the introduction of encoder backbones into advanced deep learning models with the default parameters performed very well even on the raw images. The best results were achieved using UNet++ EfficientNet-B3 (91.0%, 83.6%) and UNet++ ResNeSt (90.7%, 83.1%), establishing UNet++ as the most robust baseline when paired with powerful encoders (Fig. 5a). Interestingly, on Macenko-normalized images, the best performance was slightly lower. UNet++ EfficientNet-B3 led with 90.5% Dice and 82.8% IoU, followed closely by UNet++ Densenet121 (90.12%, 82.2%). We also noticed that some encoders, *e.g.*, DeepLabV3+ EfficientNet-B3, dropped in performance (72.3%, 57.0%), suggesting that not all architectures generalize well to stain-normalized images (Supp. Fig. S12b). The fine-tuning of parameters for UNet++ with EfficientNet-B3 on original images led to small improvements (Fig. 5b,c,d, Supp. Fig. S12c). The best performing deep learning architecture with selected parameters for Mask1 dataset was UNet++ with EfficientNet-B3 encoder model optimized with Spatial and Channel Squeeze & Excitation (SCSE) block, Softmax activation, and Cross-entropy loss function (see subsection ***Stand-alone deep learning models for semantic image segmentation task (Task 4)*** in Methods), reaching 91.6% of Dice and 84.5% of IoU values (Supp. Fig. S12c).

Similar performances were observed when using Mask2: EfficientNet-B3 with default parameters achieved 92.2% of Dice and 85.5% of IoU values, surpassing previous results from Mask1. Other well-performing encoders included EfficientNet-B4 (92.1%, 85.5%) and ResNeSt (91.2%, 83.9%) (Supp. Fig. S12d). The optimization of the same parameters as above also improved the best performing model UNet++ EfficientNet-B3, but only slightly (Supp. Fig. S10d).

Finally, we implemented and assessed three explainable AI methods, Grad-CAM (Gradient-weighted Class Activation Mapping) [76], Grad-CAM++ [77], and Saliency (Gradient-based) [78], for both raw and Macenko-preprocessed images (see subsection ***The integrated deep architecture for image segmentation and explainable AI tasks (Tasks 4 and 5)*** in Methods, Supp. Fig. S15-18). We note that it is only possible to assess the images with expert-detected and annotated ROIs, while the application of these methods can also be extended to grain-free images. To generate the saliency heatmaps, all three explainable AI methods used our best-performing MONAI+DenseNet deep learning architecture for classification. Then, we also used our best-performing optimized U-Net++/EfficientNet-B3/SCSE segmentation model from Task 4 (Fig. 1, Fig. S6a,b, Supp. Fig. S11b) to convert the saliency heat masks to binary important/non-important saliency classification masks.

For *Mask1* (Fig. 5g) using original images, Grad-CAM++ demonstrated strong performance with a Dice coefficient value of 90.7% and IoU value of 83.0%, slightly outperforming standard Grad-CAM (90.5% Dice, 82.7% IoU) and Saliency gradient-based (91.3% Dice, 84.1% IoU). The same pattern emerged for Mask1 with Macenko-normalized images (Fig. 5i), where Saliency achieved the best results (90.6% Dice, 82.9% IoU), though all methods maintained robust performance. For *Mask2*, we observed similar, while slightly improved, results. On original images (Fig. 5h), Grad-CAM++ showed marginal superiority with 91.45% Dice and 84.35% IoU compared to Saliency’s 91.23% Dice and 83.99% IoU. Interestingly, with Macenko-processed images for Mask2 (Fig. 5j), Saliency Maps performed slightly better (91.27% Dice, 84.10% IoU) than Grad-CAM++ (91.26% Dice, 83.97% IoU).

Lastly, while quantitative performance metrics (Dice, IoU) for explainability were assessed solely on images with detectable ROIs due to ground truth availability, the best-performing explainable AI method, Grad-CAM++, was also applied to grain-free images from Task 3 to generate binary heat masks, offering visual insights into the model’s decision-influencing regions in the absence of traditional pathological markers (Supp. Fig. S18).

## DISCUSSION

Here, we present a novel Skin INfectious Diseases Intelligent (SINDI) Framework, which combines shallow learning, deep learning, and explainable AI architectures to carry out tasks related to accurate diagnostics, pathogen identification, and semantic annotation of the histopathology images, assisting in decision-making and, in some cases, providing the critical information impossible to obtain by a human expert. We demonstrated the utility of our framework by applying it to one of the most devastating neglected tropical skin diseases, mycetoma. Our work is focused on targeting the most challenging tasks in skin disease diagnostics that primarily happen after the histopathology slides are obtained, and a subjective expert-based diagnosis is required. Histopathology-based diagnosis is frequently hampered by subjective interpretations and pathogen identification, leading to inter-observer variability. The latter task is especially critical due to the different treatment options required for different causative pathogenic agents. This task is also highly dependent on the quality of the histopathology slides, especially during the early onsets of the disease that are easier to treat but much harder to detect due to the lack of characteristic microbial grains. Through a comprehensive exploration of five infectious skin diagnostics challenges using 11 shallow learning, 24 deep learning, and three explainable AI strategies, three image preprocessing stages, two feature generation approaches, and various parameter optimization protocols, we provide, for the first time, a detailed assessment of machine learning feasibility for the field of infectious skin disease pathology.

Our strategy of building and assessing the SINDI framework is designed to tackle the diagnostic-related tasks in an increasing order of difficulty. In Task 1, we deal with two basic diagnostic problems: determining a disease phenotype and identifying the specific pathogen causing this phenotype. It is not surprising that the first problem, addressable by a binary classification task, is the easiest one to tackle, achieving near-perfect accuracy across both shallow and deep models and providing a strong foundation for rapid and fully automated diagnostic triage. On the other hand, pathogen identification, which is essential for guiding the treatment, presents a more challenging problem: mycetoma is known to be caused by a range of bacterial and fungal pathogens, and thus is formulated as a multi-class classification problem. For this task, the stain normalization preprocessing step, carried out by each of the three common normalization procedures used in histopathology, the Macenko, Reinhard, and Vahadane methods, showed a significant improvement. This reflects a substantial diversity in color intensity, variation, and texture that becomes critical in more challenging machine learning (ML) classification tasks. Combined with domain-driven feature generation, these two preprocessing stages are shown to be sufficient in achieving a near-perfect performance for selected shallow and deep learning models.

The questions we ask in Task 2 are already substantially more challenging for expert pathologists who, for decades, have predominantly relied on the visual identification and analysis of the regions of interest (ROIs), the areas of skin tissues affected by the pathogen and presenting disease phenotype, for diagnostics of the disease. Here, we investigate (1) how important these ROIs are for a machine learning classifier, and (2) the extent to which accurate classification can be achieved when those ROIs are occluded. We show that the ROIs are indeed important, but not critical. In fact, the four-species classification based on ROI-masked images (that we refer to as Non-ROI images) achieves a comparable performance even by a shallow ML classifier that uses Macenko preprocessing and domain-specific feature generation, the two critical steps whose importance was demonstrated in Task 1. The results from Task 2 present a paradigm-shifting advancement with significant clinical implications: it indicates that diagnostically relevant information is distributed throughout the entire image, including the regions that may appear non-informative to human experts.

Task 3 tackles what is arguably the most formidable challenge in mycetoma diagnostics: cases involving grain-free histopathology slides, *i.e.*, lacking any identifiable ROIs, due to the early stages of disease or suboptimal skin tissue sectioning. In such scenarios, definitive diagnosis typically requires time-consuming and costly molecular or microbiological assays, often inaccessible in clinics or rural hospitals. We demonstrate that advanced deep learning models can achieve accurate diagnostic performance, not only identifying each of the four pathogenic agents, but also discriminating skin tissues with disease phenotype from seven healthy skin tissue types. The success in Task 3 allows one to further advance the state of automated infectious skin disease diagnostics, reinforcing and extending the conclusions made from Task 2.

Tasks 4 and 5 shift focus from fully automated diagnostics toward enhancing interpretability and supporting pathologist decision-making. The semantic image segmentation approach in Task 4 is designed to automatically delineate pathological features, such as fungal or bacterial grains (ROIs), within the complex tissue architecture with high spatial precision. Our evaluation demonstrated the superior performance of advanced deep learning architectures coupled with optimized encoders. Notably, UNet++ employing EfficientNet-B3 encoder achieves high accuracy on both raw and

Macenko-normalized images, indicating their capacity for robust pixel-level classification of disease-relevant features. Fully automated semantic image segmentation offers an additional critical advantage: it mitigates the risk of human error in misclassifying imaging artifacts as pathological ROIs. Our method ensures that these artifacts are also highlighted as non-ROIs during the image analysis, thus enhancing the accuracy and consistency of feature identification. This task lays the foundation for downstream automated morphological analysis and feature extraction.

Complementing Task 4, the Explainable AI approach in Task 5 is focused on determining the regions, both ROIs and non-ROIs, that are critical for successful ML-based classification. By first producing the continuous saliency heatmaps and subsequently deriving simpler to interpret discrete yes/no saliency masks, this approach introduces much-needed transparency into the otherwise opaque decision-making process of deep learning models. Comparison with expert-annotated ground truths reveals strong concordance between the model’s attention regions and pathologists’ interpretation of diagnostically salient features. Furthermore, the successful application of the most accurate explainable AI architecture, Grad-CAM++, to grain-free images, despite the absence of conventional ground truth for comparison, provides invaluable insights into the model’s capacity to detect subtle, otherwise imperceptible patterns in diagnostically challenging and often unfeasible cases.

In conclusion, the SINDI framework represents a substantial and multifaceted advancement in the diagnostic landscape of infectious skin diseases, such as mycetoma. Our rigorous, five-task evaluation systematically addresses key clinical needs, consistently demonstrating the transformative potential of an integrated pipeline that combines sophisticated histopathology image preprocessing, a spectrum of machine learning methodologies (both shallow and deep learning), and transparent explainable AI techniques. At the same time, our findings open multiple promising avenues for future research and clinical translation. For universal mycetoma diagnostics, the current dataset, while comprehensive in its coverage of four predominant pathogens, could benefit from expansion to include a broader spectrum of mycetoma-causing agents and rarer clinical presentations. While our semantic feature segmentation and explainable AI methods provide valuable visual insights, future research could explore novel, perhaps more interactive, techniques tailored specifically for histopathology images to offer even more granular and intuitive explanations to pathologists, potentially incorporating attention mechanisms or counterfactual explanations. Lastly, we envision that the ultimate test for the SINDI framework will be its performance in diverse, real-world clinical settings. Careful consideration of the practical integration of such an AI tool into existing pathology workflows is crucial. This includes addressing infrastructure requirements, user interface design for seamless interaction, and robust ethical considerations surrounding AI-assisted diagnosis. By addressing the current limitations and adapting the SINDI framework to the diagnostics of other infectious skin diseases, we hope to achieve its full potential in revolutionizing ML-assisted diagnosis and ultimately improving patient care globally.

## METHODS

The goal of this work is to develop an advanced machine learning approach that integrates cutting-edge deep learning and explainable AI paradigms to understand the critical factors that influence the diagnosis of infectious skin diseases across the diverse bacterial or fungal pathogens responsible for these diseases, with the aim of developing a robust computational diagnostic tool. This tool would not only assist pathologists in accurate species identification but would also provide transparent insights into the decision-making process behind the diagnosis. Here, we focus on one of the major tropical skin diseases, mycetoma, introducing a novel machine learning approach, the Skin INfectious Disease Intelligent (SINDI) Framework. The SINDI framework offers an automated, comprehensive, and interpretable solution for mycetoma diagnosis. Specifically, our approach seeks to address five pivotal questions that pathologists encounter in clinical practice. First, how can one accurately diagnose the four most recurrent mycetoma species in Senegal and other countries of the “Mycetoma Belt” [13] and distinguish between eumycetoma (mycetoma caused by fungal species) and actinomycetoma (mycetoma caused by bacterial species)? Second, can an accurate diagnosis be achieved even when histopathology images lack clearly defined regions of interest (ROIs), the biological markers of the infections, and how does the presence of either fully defined or masked ROIs in the histopathology images impact the diagnostic accuracy and clinical utility of the model? The ROIs, also referred to as “grains” are fine sand-like particles that are small clusters of pathogens surrounded by an inflammatory reaction and stained in the images. Third, can an automated method address an even more challenging case, where the histopathology image completely lacks any ROIs (grain-free)? Fourth, can one automatically determine ROIs in the histopathology images, assisting the clinicians in diagnosis and treatment planning? Lastly, can an automated approach provide clinicians and pathologists with transparent insights into the model’s decision-making process, fostering trust and understanding?

In addressing these questions, we introduce a novel deep learning and explainable AI framework that balances high diagnostic accuracy with interpretability. Our SINDI framework includes seven core components (Fig. 1) to ensure a comprehensive approach to mycetoma diagnosis. First, the *Study Population and Data Collection* component involves gathering, manual curation, and annotation of a diverse dataset of histopathology images, representing infections by the most recurrent mycetoma species. Second, the *Image Preprocessing and Normalization* component addresses variability in staining through advanced imaging methods, such as Macenko [64], Reinhard [71], and Vahadane [68]. Third, the *Image Annotation* component leverages expert-based manual annotation to establish ground truth masks. Fourth, in the *Supervised Learning Task* component, we prepare the datasets for the assessment on four different classification tasks: (1) disease and species classification of the images with ROI present (Task 1); (2) disease and species classification of the images with ROI masked (Task 2); (3) disease and species classification of the images without any ROIs present (grain-free, Task 3); and (4) semantic image segmentation to isolate ROIs in the image (Task 4). To do so, datasets are generated that include ROIs, Non-ROIs, and grain-free scenarios. Fifth, the *Supervised Shallow and Deep Learning for Classification and Segmentation* component trained and evaluated both the standard shallow learning as well as advanced deep learning models, including DenseNet-121 with MONAI and U-Net++ with EfficientNet-B3 models, to achieve species classification and automated semantic segmentation. Sixth, the *Feature Engineering and Analysis* component focuses on extracting the texture, color, and morphological features and utilizing feature embedding techniques, such as t-SNE and UMAP, for data visualization and structure analysis. Last, the *Explainable AI (XAI)* component (Task 5) integrates Grad-CAM and Grad-CAM++ methods as well as saliency maps to provide interpretable visualizations of the model’s decision-making process. Together, these seven components form a cohesive framework for developing an accurate, transparent, and clinically valuable diagnostic tool for mycetoma.

### Task 1: Identification of disease samples and the source pathogens

The first task is the most straight-forward and is designed to simulate the diagnostics scenario a pathologist is faced with (Fig. 2a). Specifically, for a give histopathology sample of a disease patient, our method determines (1) if the sample comes from the disease or healthy patient (disease classification task), and (2) if it comes from the disease patient, the source pathogen causing the disease (species classification task). While the first subtask is formulated as a standard binary classifier problem, the second subtask presents a more challenging *N*-class classification problem, where *N* is the number of pathogenic species considered. In our case, we considered four major pathogens (see Study Population and Data Collection subsection for more details). For this task, we use an original dataset of raw histopathology images, as well as three additional datasets generated through stain normalization techniques: Macenko [64], Vahadane [68], and Reinhard [71]. Each image serves as a source of numerical features. First, a domain-independent feature extraction strategy is employed with the Img2Vec [79] package, which utilizes a pre-trained ResNet [80] network to extract 512-dimensional feature vectors. These feature vectors serve as the input for a range of machine learning classifiers. Second, a domain-specific feature extraction strategy is employed as an alternative approach. We extract the features that are categorized into several domains relevant to histopathology imaging, such as morphology, color and intensity, texture, statistics, as well as structural and spatial features (see

Feature engineering subsection for more details). To visualize the relationships between mycetoma species within the high-dimensional domain-specific feature spaces, we use two embedding techniques, UMAP (Uniform Manifold Approximation and Projection) [81] and t-SNE (t-Distributed Stochastic Neighbor Embedding)[82] (Fig. 1, Fig. 2g, Supp. Fig. S4, Supp. Fig. S6). These techniques allow us to visualize substantial improvement in class separability when using preprocessing steps compared to the original feature space.

For a comprehensive assessment of supervised learning models’ performance on the two subtasks of Task 1, 3 machine learning classifiers, commonly used for multiclass image classification, are trained for the input set of domain-independent feature vectors. The classifiers include decision tree [83], random forest [84], and logistic regression (linear kernel) [85] (Fig. 2b,d, Supp. Fig. S3a, Supp. Fig. S5a, Supp. Fig. S7). Furthermore, 10 classifiers (Fig. 2e,f, Supp. Fig. S3b, Supp. Fig. S5b, Supp. Fig. S6) are trained for the input set of feature vectors generated through a domain-specific selection strategy, including linear (logistic regression, Support Vector Machines (SVMs) with linear kernel, and Naive Bayes), non-linear (K-nearest neighbors (KNN), decision trees, Random Forests, gradient boosting, CatBoost, XGBoost [86], and Quadratic Discriminant Analysis (QDA).

### Task 2: Assessment of ROI and Non-ROI impact on species classification accuracy

During the diagnostics of infectious skin diseases, pathologists primarily rely on detecting and characterizing specific areas of histopathology images, known as regions of interest (ROIs), that inform the disease phenotype. However, the quality of the ROIs (also referred to as “grains”) in an image varies drastically and could be a source for misdiagnosis or misidentified pathogen species. Thus, in the second task, we test the ability of our approach to correctly classify pathogenic species even when the ROIs are completely “scrambled” and carry no information, rather than their contours. To do so, we first manually annotate each histopathology image by isolating ROI and Non-ROI areas, and then generating two groups of images: (1) ROI images, where all pixels except the ones belonging to ROIs are replaced with the same average value; and (2) Non-ROI images, where all pixels belonging to ROIs are replaced with the average value. Finally, we assess the contributions of ROIs to the species classification by comparing the accuracy of the same supervised learning algorithms independently trained on ROI *vs.* Non-ROI datasets (Fig. 1, Fig. 3a, Supp. Fig. S8a).

Because of the superior performance of Macenko normalization, here we consider images with this preprocessing step as a source of non-ROI and ROI images. We then compare the classification performances on those two datasets to those based on the Non-ROI and ROI datasets generated from the original, raw images. From each of the four datasets, we extract a comprehensive set of domain-specific features (Supp. Fig. S8c). The feature set includes color, morphological, texture, and statistical characteristics. To visualize the impact of isolating ROI and Non-ROI based features on the separability of the mycetoma species, we again use UMAP [81] and t-SNE [82] embeddings (Fig. 1, Fig. 3d,e, Supp. Fig. S8d). For each of the four datasets, we train and assess the same series of 10 shallow learning classifiers as well as MONAI-DenseNet121 [87] deep learning approach embeddings (Fig. 1, Fig. 3b,c, Supp. Fig. S8b,c) as in Task 1.

### Task 3: Species classification for images with no visible grains

Next, we test the capacity of a supervised learning approach to perform diagnostics in cases that are difficult or even impossible for a pathologist. Specifically, for this task, our methods are trained on a dataset of grain-free images, where no ROI can be detected. The task is expected to be extremely challenging not only for a human pathologist but also for a supervised classifier. To conduct this analysis, we utilize two distinct datasets: a generated grain-free dataset, consisting of image segments lacking any visible ROIs, and a combined dataset, which integrated the grain-free images with the ‘Skin Tumor Histology Dataset’, serving as a negative control (see ***Generation of grain-free datasets*** subsection for more details) (Fig. 1, Fig. 4a, Supp. Fig. S9a). From these preprocessed datasets, we extract a similar range of features as in Tasks 1 and 2: color, morphology, texture, intensity, and statistical measures (Fig. 4f, Supp. Fig. S9). We then employ the same series of 10 shallow learning classifiers as well as the MONAI-DenseNet121 deep learning approach (Fig. 4b,c,d,e, Supp. Fig. S9b,c). The classifiers are trained and evaluated separately on both the grain-free datasets and the combined dataset. This allows us to assess the classifiers’ ability to distinguish between mycetoma species in the absence of ROIs, and also to evaluate their ability to distinguish between mycetoma species and various types of healthy skin tissues.

### Task 4: Deep learning approaches to semantic image segmentation

Next, we develop an automated image segmentation model using convolutional neural networks (CNNs) for precise semantic segmentation. This model classifies each pixel into predefined categories, segmenting the image into meaningful regions, which correspond to distinct structures or tissues. In the context of mycetoma, the segmentation is critical for differentiating between mycetoma species and healthy tissues, ultimately facilitating faster and more accurate diagnostics by a pathologist. Our semantic segmentation model consists of two key components: the encoder (feature extractor) and the decoder (reconstructor). The encoder progressively reduces the image size (downsampling) while extracting essential features. The decoder, on the other hand, upsamples the image to the original resolution, filling in the details of the segmentation mask. For this task, we again use Macenko-preprocessed and original images as two independent sources. We then perform two-expert manual annotations of the images, resulting in two distinct sets of ground truth masks representing ROIs being generated for the same image dataset (see ***Manual segmentation for ground truth establishment*** subsection for more detail). Lastly, we implement and compare 12 deep learning model architectures (Fig. 1, Supp. Fig. S11a, see ***Stand-alone deep learning models for semantic image segmentation tasks*** and ***The integrated deep architecture for image segmentation and explainable AI tasks*** subsections for more details).

### Task 5: Leveraging Explainable AI to develop clinical guidelines for classification and segmentation

As our last task, we developed an explainable AI approach to guide the diagnostic decision-making by a pathologist. Specifically, we use three independent methods, Grad-CAM (Gradient-weighted Class Activation Mapping) [76], Grad-CAM++ [77], and Saliency (Gradient-based) [78] to address two subtasks. First, we automatically determine the more and less important regions for the classifier’s decision-making by generating saliency heatmaps (Fig. 1 and Suppl. Fig. S1a). The saliency heatmap is based on a continuous measure that quantifies the contribution of different parts of the input image to the classification task. Second, we integrate the generated saliency heatmaps with the corresponding original images (both raw and Macenko-preprocessed images) as a modified input for a deep learning classifier to divide the continuous-values heatmap into a binary important/non-important saliency classification mask. In contrast to the continuous-values saliency heatmap, the saliency classification mask is a binary mask, thus providing the pathologist with a simple additional layer of information about the important parts of the histopathology image making the diagnosis more informed (Fig. 1, Fig. S6a,b, Supp. Fig. S11b, see ***Explainable AI for histopathology images with detectable ROIs*** and ***Explainable AI for grain-free images*** subsections for more detail). Importantly, the explainable AI protocol was applied and tested for both images with the detectable ROIs as well as grain-free images.

### Study population and data collection

For this study, we collect the histopathology imaging and clinical data from a Senegalese population diagnosed with mycetoma, extending our initial dataset [21] from 1,289 to 1,324 images. The dataset includes samples from 34 patients, comprising 71 histopathology slides, with each patient contributing between 1 and 6 slides (Fig. 1, Fig. 2c). It represents the four most common mycetoma-causing species in Senegal, which are also widely spread in other countries of the “Mycetoma Belt” [13]: *Actinomadura pelletieri* (AP), *Falciformispora senegalensis* (FS), *Madurella mycetomatis* (MM), and *Streptomyces somaliensis* (SS).

Histopathology images (Fig. 1 – DATA COLLECTION, Fig. 2a) are acquired using a Leica ICC50 E microscope at three magnification levels: 4x, 10x, and 100x. Tissue samples are stained with Hematoxylin and Eosin (H&E), a widely used staining technique in histopathology that highlights cellular and tissue structures [26]. This staining facilitates the visualization of key tissue features, enhancing the presence of grains within the samples. Images are captured in .jpg format with dimensions of 1200×1600 pixels and a resolution of 24 bits per pixel color. Each image is then used to establish a diagnosis by two experienced pathologists, ensuring accurate species identification.

### Data preprocessing and normalization

During the initial analysis, we observed significant color variability across histopathology images, even among samples of the same species from different patients. This variability, often caused by differences in staining protocols, scanners, or acquisition settings [88], [89], can confound machine learning models and hinder accurate species identification. To address this, we applied and compared three stain normalization techniques[63] to ensure image consistency across all datasets. Specifically, we employ the Macenko [64], Reinhard [71], and Vahadane [68] methods that were originally developed to process histopathology images of other diseases and tissues (Fig. 1-IMAGE PREPROCESSING). Macenko method [64], known for its flexibility in handling staining variations, transforms the color space to match a reference stain, typically Hematoxylin and Eosin (H&E). Its consistency may be affected when applied to the images with complex staining variations. Reinhard [71] normalizes color distributions by aligning the target image to a reference, offering computational efficiency and simplicity. It can perform well when the reference and target images share similar characteristics, but may degrade with large staining variations. Vahadane [68] utilizes color deconvolution to separate stain components, making it particularly effective for datasets with significant staining differences and preserving tissue features well, albeit with a higher computational cost. By generating three additional datasets using these normalization methods alongside the original dataset, we improved consistency and reliability, facilitating robust analyses and ensuring the reproducibility of our results.

#### Macenko normalization

The Macenko normalization method [64] focuses on normalizing histology slides and tackles the inherent variability in staining through a two-stage process. First, it corrects stain vector variation by converting initial RGB pixel values to optical density (OD) space and subsequently removes pixels with an overall OD below a threshold (β=0.15). Singular Value Decomposition (SVD) is then applied to the remaining OD data, and the pixels are projected onto a plane defined by the top two singular vectors. The angle of each projected point is calculated, and robust extremes are identified and converted back to OD space to determine the optimal stain vectors for the specific slide. These vectors are later used in a color deconvolution scheme to separate the stain contributions. Second, to correct intensity variations, pixels dominated by each stain are identified, and their intensity distributions are characterized by histograms. A robust pseudo-maximum intensity is determined for each stain by calculating the 99th percentile of its intensity histogram. Finally, all intensity histograms for a given stain across different slides are scaled to match this pseudo-maximum, normalizing the overall stain intensity and enabling more reliable quantitative comparisons.

#### Reinhard normalization

The Reinhard normalization method [71] focuses on color normalization and involves transforming the color distribution of a target image to match that of a reference image. Their approach involves converting both a source and a target image into the decorrelated lαβ color space, a space designed to minimize channel correlations and better align with human perception. Once in this space, the mean and standard deviation of each color channel (l for achromatic channel, α for chromatic yellow-blue, and β for chromatic red-green) are calculated for both images. The color characteristics of the source image are then imposed onto the target image by adjusting the mean and standard deviation of the target image’s lαβ channels to match those of the source. Finally, the modified target image is transformed back from the lαβ color space to the standard RGB color space for display.

#### Vahadane normalization

The Vahadane normalization method [68] was developed for structure-preserving color normalization (SPCN). It is designed to address color variations in histopathology images caused by factors such as staining techniques and scanner differences. The approach involves two key steps: (1) Stain Separation using Sparse Non-negative Matrix Factorization (SNMF) and (2) Color Normalization based on the estimated stain densities. In SNMF, the input image is first transformed into optical density using the Beer-Lambert law. The core of SNMF is to decompose the image into a non-negative matrix factorization, which incorporates a sparsity constraint to ensure that each pixel represents a sparse mixture of biological materials. For the color normalization step, one uses the stain density map from the source image and combines it with the color basis from a target image, thus preserving the biological structure of the source while altering only its color appearance. This technique ensures that only the color appearance changes, preserving structural integrity. Finally, to accelerate the computation on large whole-slide images (WSI), a patch-based sampling strategy is used to estimate the color appearance matrix for the image, which is then applied to smaller patches of the WSI, reducing computational cost.

#### Manual segmentation for ground truth establishment

To establish ground truth for training our segmentation models, we carry out manual annotation of ROIs across all 1,324 histopathology images. The annotation is focused on the precise delineation of regions containing mycetoma grains, which are critical for a diagnosis by a human specialist [26]. Utilizing Label Studio (version 1.13.1) [90], each image is independently reviewed and annotated by two expert pathologists (Fig. 1-IMAGE ANNOTATION). This two-expert annotation strategy is implemented to ensure the accuracy and inter-observer consistency, mitigating potential biases and variations in interpretation. As a result, two distinct sets of ground truth masks are generated for the same image dataset, which we designate as *mask1* and *mask2*. The mask1 and mask2 datasets are then used and comparatively assessed in our Image Segmentation and Explainable AI tasks. The creation of these high-quality, pathologist-validated ground truth datasets is critical for training robust segmentation models, serving as the definitive benchmark for learning and subsequent evaluation.

#### Region of Interest (ROI) and Non-ROI Dataset Generation

To investigate the relevance of different image regions for species classification, we perform ROI/Non-ROI image partitioning. This procedure involves utilizing the original histopathology images and their corresponding ground truth segmentation masks to create two distinct image sets. Each original image is divided into two complementary images (Fig. 3a, Supp. Fig. S6a): (1) the ROI image, which retains only the pixel values corresponding to the annotated regions within the mask, and (2) the non-ROI image, which contains the pixel values outside these regions. We note that this procedure does not affect the size of the image: to isolate the non-ROI areas, the real pixel values within these regions are replaced with the pixels of an average color of the original image. This process was applied to both the original dataset and the Macenko-normalized dataset, resulting in a total of four datasets: ROI and non-ROI sets for each. The Macenko normalization was selected because of its superior performance on the original classification task (Fig. 2e,f).

The average color, *C_avg_*, of the original image ***I*** with dimensions *H×W* is defined as follows:

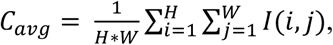

where *I(i,j)* represents the pixel value at position *(i,j), a*nd the operation is performed across RGB channels.

The masked images of the two types, *I_ROI_* and *I_Non-ROI_*, are then defined using the mask *M*, where for a pixel (*i*,*j*), *M*(*i*,*j*) > 0 indicates that this pixel belongs to an ROI. Specifically:

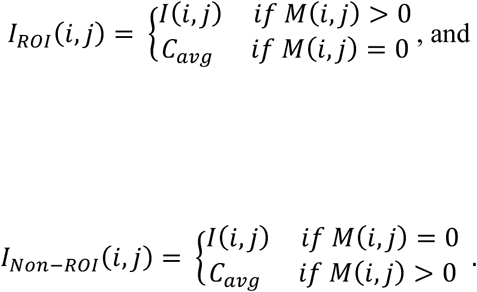

#### Generation of grain-free datasets

To investigate the feasibility of species classification in the images that completely lack any visible ROIs (grains) and therefore would be nearly impossible to diagnose by a pathologist without any follow-up experimental study, we generate a dataset of grain-free images. To do so, a systematic preprocessing approach is implemented, consisting of two basic steps. First, each image from the original dataset is divided into eight equal-sized image fragments. Second, the fragments containing any portion of the annotated ROI, indicative of grain presence, are discarded, leaving only those fragments that are free of ROIs. This procedure is also applied to the Macenko normalized dataset. As a result, two distinct grain-free datasets are generated (Fig. 4a, Suppl. Fig. S7a). Additionally, to establish robust negative control, comparing the disease grain-free samples with the healthy samples of different skin tissues, the Skin Tumor Histology Dataset [54] is integrated into our analysis. The latter dataset contains images of tumor and non-tumor tissue types; for this work, only non-tumor classes representing healthy tissues are considered. Since mycetoma primarily affects subcutaneous tissue, the inclusion of healthy tissue from subcutaneous layers, such as subcutis, is particularly relevant. The dataset also includes other tissue types associated with skin, including epidermis, dermis, sebaceous glands, nerves, skeletal muscle, and vessels, providing a diverse range of healthy tissue types that can theoretically strengthen the model’s ability to generalize well. For computational efficiency, 1,000 images are extracted from each of the selected classes (seven classes from the “Skin Tumor Histology Dataset”), resulting in a total of 7,000 images (Fig. 4a). With the images of healthy tissues integrated, our final dataset includes 11 classes: four mycetoma species (3,034 images) and seven healthy tissue types (7,000 images).

### Supervised learning approaches for disease and species classification tasks (Tasks 1-3)

For Tasks 1-3, shallow and deep learning supervised classifiers are developed using vectors of features extracted via two distinct strategies: domain-independent and domain-specific extraction. The domain-independent features are then fed into three types of shallow learning classifiers applied to Task 1, including decision trees, random forests, and logistic regression. This feature extraction strategy is not considered for more advanced classification models as well as in Tasks 2 and 3 due to its inferior performance when applied in Task 1. Logistic regression, further extended for the multiclass (pathogenic species) classification via one-vs-rest, or softmax, regression, is evaluated with six different solvers: *lbfgs*, *liblinear*, *newton-cg*, *sag*, *saga*, and *saga l2*. Decision trees are selected for their interpretability, since they recursively partition the feature space. To mitigate potential overfitting, hyperparameters, including maximum tree depth and minimum samples per split, are optimized. For the random forests, the optimized hyperparameters include the number of trees, maximum depth, and feature subset size. For the parameter optimization, we implement both the parameter grid search (exhaustive hyperparameter exploration) and the randomized search (efficient sampling from parameter distributions).

The dataset of feature vectors extracted with the domain-specific strategy is used to train 10 shallow learning and one deep learning supervised classifiers grouped into linear (logistic regression, Support Vector Machines (SVMs) with linear kernel, and Naive Bayes), non-linear (K-nearest neighbors (KNN), decision trees, Random Forest, gradient boosting, CatBoost, XGBoost, and Quadratic Discriminant Analysis (QDA). The classifiers are initialized with specific hyperparameters to optimize performance. For Logistic Regression, the training involved a maximum of 200 iterations, a One-vs-Rest multi-class approach, and the liblinear solver. The SVM, configured as a OneVsRestClassifier with SVC, uses a linear kernel while enabling probability estimates. Gaussian Naive Bayes is implemented under the default probabilistic parameters. The KNN model considers the four nearest neighbors for predictions. Decision tree implementation relies solely on a predefined random state. The Random Forest incorporates 100 estimators and a fixed random state for consistency. The Gradient Boosting classifier employs 100 estimators and a fixed random seed. CatBoost runs 1000 iterations with a learning rate of 0.1, a tree depth of 6, and periodic training logs. XGBoost is implemented with a multinomial logistic loss as the evaluation metric, a softmax objective for multi-class classification, and an automatically determined number of classes from the label encoder. Lastly, QDA is implemented using the default settings, including standard quadratic decision boundaries.

For the deep learning architecture, we apply the Medical Open Network for AI (MONAI) framework [87] with a pre-trained DenseNet121 backbone [91]. Here, we rely on the model’s intrinsic ability to extract the features instead of employing the above two feature extraction strategies. MONAI’s medical imaging-specific optimization, such as efficient patch-based inference and standardized preprocessing, combined with DenseNet121’s ability to hierarchically learn discriminative features, make this architecture a good fit for our tasks.

### Stand-alone deep learning models for semantic image segmentation task (Task 4)

Using the Macenko and original dataset, and corresponding ground truth, we implement 12 deep learning model architectures (Fig. 1), including UNet++ [92], UNet [93], LinkNet [94], SegFormer [95], FPN (Feature Pyramid Networks) [96], DeepLabV3 [97], DeepLabV3++ [98], PAN (Path Aggregation Network) [99], SegNet [100], Attention UNet [101], MANet (Multi-scale Attention Net) [102], and PSPNet (Pyramid Scene Parsing Network) [103]. After testing models using default encoders, we then expand our approach by incorporating pre-trained backbones using the segmentation-models-pytorch library [104]. Specifically, we replace the encoder part of the semantic segmentation models with various pre-trained models, including EfficientNet [105], DenseNet [91], ResNet [80], MobileNet [106], VGG [107], Inception [108], Xception [109], SE-Net (Squeeze-and-Excitation Networks) [110], DPN (Detail-Preserving Network) [111], ResNeSt (Split-Attention Networks) [112], Convolutional feature extractor, and Mix vision transformer to improve convergence and performance. In total, we explored 40 architecture-encoder combinations. Lastly, when for a given encoder there are multiple available versions (*e.g.*, efficientnet-b0, efficientnet-b1, …, efficientnet-b7), we select the one based on its performance on the mask1 original set (see ***Manual segmentation for ground truth establishment*** subsection).

Once the evaluation is done (see ***The integrated deep architecture for image segmentation and explainable AI tasks (Tasks 4 and 5)*** subsection), we select the best-performing, in terms of dice coefficient and IoU value, combination of architecture-encoder—UNet++ with EfficientNet-B3—and attempt to improve it further through several optimization strategies. First, data augmentation techniques [113], such as rotation, flipping, and color jittering, are employed to improve model robustness and mitigate overfitting. Second, learning rate scheduling[114] is implemented using (1) StepLR scheduler [115], which dynamically adjusts the learning rate during training by reducing it by a factor of gamma every step_size epoch, (2) ReduceLROnPlateau [115], which reduces the learning rate when the validation loss plateaus, and (3) CosineAnnealingLR [115], which uses a cosine annealing schedule to adjust the learning rate (Supp. Fig. S10e). Third, we also explore the Dual Encoder UNet++ model [116], which extends the original model by combining EfficientNet-B3 and DenseNet-121 encoders within the same architecture, leveraging the strengths of both encoders for better segmentation results. Fourth, we add Spatial and Channel Squeeze & Excitation (SCSE) blocks [117] to enhance relevant features and suppress noise, improving boundary accuracy and overall segmentation quality. Fifth, to further optimize training dynamics [104], [118], we evaluate two adaptive optimizers: Adam [115], which combines momentum and RMSProp for efficient convergence, and AdamW [115], an alternative version with decoupled weight decay to improve generalization. For the decoder architecture, we experiment with channel configurations [(384, 192, 96, 48, 24), (512, 256, 128, 64, 32), (1024, 512, 256, 128, 64), (256, 128, 64, 32, 16)] to balance the feature extraction efficiency and computational cost. The pooling strategy [118] is tested with both average (“avg”) and max pooling (“max”) operations, while decoder interpolation [118] modes—“nearest” (default), “bilinear”, “bicubic”, and “nearest-exact“—are compared to refine the upsampling precision. We then assess Softmax (for interpretable probabilities) and LogSoftmax (paired with NLLLoss for stable gradients) as activation functions, ensuring robustness across different training scenarios [118]. As a last optimization strategy, we explore three different loss functions during the model (UNet++ with EfficientNet-B3) training: Cross-Entropy Loss [118], a combination of Cross-Entropy Loss and Dice Loss[118], and Focal Loss [118].

### The integrated deep architecture for image segmentation and explainable AI tasks (Tasks 4 and 5)

We next extend standard deep learning architectures to address more challenging image segmentation and explainable AI tasks. While a stand-alone classifier may suffice for standard classification tasks, integrating multiple deep demonstrated architectures can offer substantial advantages in more complex settings. Previous studies have demonstrated improvement in the image segmentation task for breast cancer histopathology images when combining UNet and EfficientNet architectures. We further extend this approach to image segmentation of tropical skin diseases by integrating three different architectures: we combine U-Net++ [92] with EfficientNet-B3 [105] that replaces the default encoder of U-Net++, and SCSE [117] that enhances the U-Net++ default decoder module (Supp. Fig. S9a). After evaluating the performance of the integrated U-Net++/EfficientNet-B3/SCSE architecture [104], [118] and demonstrating its superior performance compared to the stand-alone deep learning architectures, we use the same integrated architecture for the explainable AI tasks (see Subsection

#### Explainable AI for histopathology images with detectable ROIs and Explainable AI for grain-free images)

One of the main reasons for selecting the U-Net++ architecture is that it refines the traditional U-Net by introducing nested, densely connected skip pathways between encoder and decoder layers. This design bridges semantic gaps in feature maps, improving gradient flow and hierarchical feature integration. The skip pathways link intermediate encoder layers to their corresponding decoder layers, enriching the decoder with multi-resolution information. Each feature map, *x^i,j^*, is computed as:

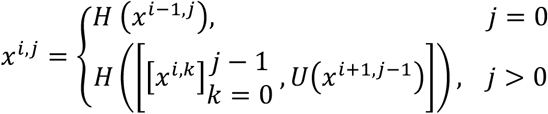

where *H(⋅)* denotes convolution followed by activation, *U*(*⋅*) is upsampling, and [*⋅*] indicates concatenation.

In the integrated U-Net++/EfficientNet-B3/SCSE architecture, the encoder, tasked with multi-scale feature extraction, employs EfficientNet-B3 due to its optimal balance of computational efficiency and accuracy. EfficientNet-B3 comprises seven Mobile Reversed Bottleneck Convolution (MBConv) blocks, progressively downscaling the input image size from 224×224 to 7×7, while extracting intermediate feature maps sized 112×112, 56×56, 28×28, and 14×14. These maps are routed to the decoder via U-Net++’s skip pathways, enabling precise segmentation of fine-grained structures. To further optimize feature recalibration, SCSE blocks are then embedded in the U-Net++’s decoder. These blocks employ dual mechanisms: (1) Channel Squeeze and Excitation, which compresses spatial information into channel descriptors via global average pooling, then scales channels by learned weights; and (2) Spatial Squeeze and Excitation, which uses convolutional attention maps to highlight critical spatial regions. Together, these mechanisms amplify relevant features while suppressing noise, improving segmentation of overlapping or subtle structures.

### Feature engineering and visualization

The inherent complexity and heterogeneity of histopathology images complicate the distinction between mycetoma species, demanding a comprehensive feature extraction process. This process aimed to quantify relevant image characteristics, establishing a robust foundation for automated classification that surpassed subjective visual assessments [119], [120], [121], [122], [123], [124]. The feature engineering for this work includes two basic strategies: domain-independent and domain-specific. For the first approach, we utilized the ImgVec package [79] to extract 512-dimensional feature vectors from histopathology images using a pre-trained ResNet network [80]. For the second approach, we extracted domain-specific features tailored to histopathology, including morphology, color/intensity, texture, statistical, and structural/spatial attributes. Morphological features such as area, perimeter, and circularity are calculated, with the circularity measure defined as:

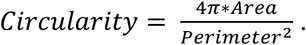

These features provide quantitative measures of grain and tissue shape, often indicative of species-specific characteristics. Color and intensity features, such as mean intensity, median intensity, intensity variance, and mean RGB values, provide quantitative measures of tissue chromatic properties, capturing subtle color variations that may distinguish species. Texture features, derived from the Gray-Level Co-occurrence Matrix (GLCM), including contrast, correlation, and energy, and local binary pattern histograms, capture spatial relationships and patterns in the grayscale images, revealing subtle textural differences that may reflect underlying tissue structure and species-specific patterns. Statistical features, including mean intensity, standard deviation of intensity, skewness, and kurtosis, describe the intensity distribution within the images, highlighting variations in tissue homogeneity and heterogeneity, which can be indicative of different pathological processes. Lastly, structural and spatial features, such as cell density and perimeter density, represent the spatial arrangement of tissue components, providing insights into tissue organization and the distribution of grains within the tissue.

To visualize the high-dimensional feature space and explore complex relationships between mycetoma species and the healthy tissues, we employ two standard non-linear dimensionality reduction methods: t-SNE (t-Distributed Stochastic Neighbor Embedding) [82]and UMAP (Uniform Manifold Approximation and Projection) [81]. These techniques are essential for understanding the underlying structure of the data and identifying potential patterns that could aid in species classification. The results can also potentially highlight the advantage of using some preprocessing methods over the others. The two methods are applied to visualize the feature vectors generated using a more accurate domain-specific strategy for both the raw images and the images preprocessed using the three methods (Macenko, Reinhard, and Vahadane).

### Explainable AI for histopathology images with detectable ROIs

The two-subtask protocol to incorporate Explainable AI [125], [126], [127], [128] information to images with detectable ROIs, consists of: (1) saliency heatmaps generation and (2) integration of the heatmaps into the semantic segmentation task (originally carried out in Task 4). The protocol’s first subtask leverages three methods: Grad-CAM (Gradient-weighted Class Activation Mapping) [76], Grad-CAM++ [77], and Saliency [78] (Fig. 1). The methods, in turn, are based on the MONAI-DenseNet121 model originally trained for four-class classification (Task 1). Grad-CAM (Gradient-weighted Class Activation Mapping) produces continuous-value saliency heatmaps that identify regions in the image contributing to a specific prediction (Fig. 6a,b). It calculates gradients of the class score with respect to feature maps of the last convolutional layer, averages these gradients, and applies a ReLU activation to focus on positively contributing regions. Grad-CAM++ extends Grad-CAM, incorporating higher-order derivatives to generate sharper and more accurate saliency heatmaps, with particular improvements when the image contains multiple objects or fine-grained details. The last method, Saliency Maps (Gradient-based), computes the gradient of the output class score with respect to each input pixel, revealing the most influential pixels for a given prediction. The magnitude of these gradients indicates the importance of each pixel, resulting in fine-grained heatmaps.

**Figure 6.**
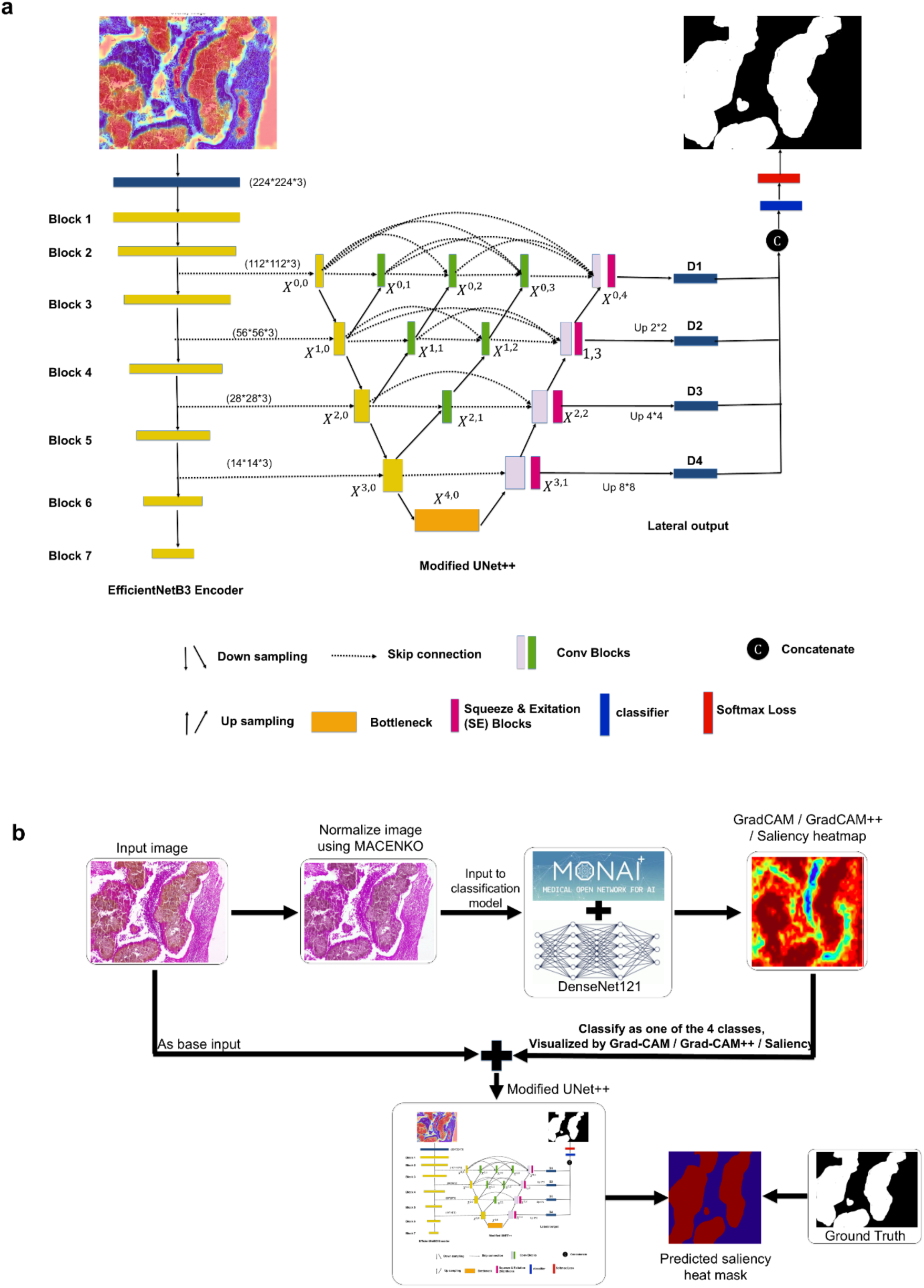
Best-performing semantic segmentation architecture and explainable AI workflow. **(a) Optimized Semantic Segmentation Architecture.** The architecture of the best-performing semantic segmentation model, UNet++ with an EfficientNet-B3 encoder and SCSE (Squeeze-and-Excitation) blocks. The same architecture is leveraged for the explainable AI workflow **(b)**, with the key distinction of accepting a six-channel input image (original image combined with the saliency heatmap image) for explainable AI task, compared to three channels for semantic segmentation (original image). The overall explainable AI workflow includes the process and key steps involved in Task 5, from input processing to the generation of interpretable outputs for clinical guidance.

For the saliency heatmaps generation, we apply Grad-CAM, Grad-CAM++, and Saliency (Gradient-based) methods to the raw images as well as images preprocessed using the top-performing in Task 1 Macenko method. For both Grad-CAM and Grad-CAM++, the last convolutional layer of the DenseNet-121 model (prior to the fully connected layer) serves as the target layer for gradient calculation. For Grad-CAM and Grad-CAM++, we utilized the MONAI framework’s visualization library, targeting the ReLU activation layer to maximize spatial and interpretative details. Saliency maps are computed using a custom function that performs forward and backward passes to calculate gradients and highlight influential pixels.

For the second subtask, we integrate the obtained saliency heatmaps with the raw images as well as the Macenko-preprocessed ones. As a result, the input image has six channels instead of the original three. We then retrain a U-Net++/EfficientNet-B3/SCSE architecture that is originally used for the segmentation task (Task 4). As in the previous cases, we use the ground-truth masks from the manual annotations to guide the segmentation process, minimizing the loss function. The generated saliency heatmap from the original and Macenko-normalized datasets using Grad-CAM, Grad-CAM++, and Saliency maps form six additional datasets in total (Fig. 6a,b). The output of this procedure is a binary saliency classification mask, providing a pathologist with a streamlined way to convert the complex continuous-value saliency heatmap information to a much simpler yes/no mask.

### Explainable AI for grain-free images

To help pathologists understand the results of the most challenging grain-free image classification task (Task 3), we apply a similar two-subtask explainable AI protocol modified to process the grain-free images. For the first subtask, the grain-free saliency heatmaps are generated using Grad-CAM++. This method is selected due to its superior performance on the previously mentioned explainable AI task (images with detectable ROIs), compared to the other two methods. For the second subtask, to integrate the saliency heatmaps into the semantic segmentation workflow, they are combined with the original images (raw and Macenko-processed) as an input for the semantic segmentation deep learning model (the same U-Net++/EfficientNet-B3/SCSE architecture as in Task 4 is used). As a result, the saliency classification masks are produced that emphasize the important areas that are hard or impossible to detect by a human expert, providing an additional information layer for the decision making (Sup Fig. 9b).

### Evaluation protocol

The evaluation of each classification and explainable AI task is carried out individually, independently of each other, and using different datasets for the ground truth. For Task 1, we randomly split the manually labeled dataset into 70% (926 images), 10% (132 images), and 20% (266 images) for the training, parameter optimization, and testing stages, respectively. In order to limit the information leakage between the stages, we ensured that the samples from each patient were exclusively in one of three subsets. For both subtasks, healthy (7000 images) vs. disease (1324 images) and 4-class pathogen classification (*Madurella mycetomatis, Falciformispora senegalensis, Actinomadura pelletieri, and Streptomyces somaliensis*), four evaluation criteria are calculated: accuracy (*Acc*), precision (*Pr*), recall (*Re*), and F1-score (*F*1):

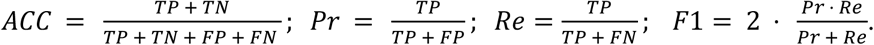

Here, for the healthy vs. disease binary classification, True Positives (*TP*), True Negatives (*TN*), False Positives (*FP*), and False Negatives (*FN*) are calculated with disease being the positive class and healthy being the negative class. For the 4-class pathogen classification, *TP* is the number of all correctly identified class members across all four classes, and *TN* is the number of all correctly identified class non-members across all four classes. *FP* is calculated for the individual class using a one-*vs*-all scenario (*i.e.*, for the first class, the members from all other classes that are misidentified as the first class members are considered false positives for this class), and these *FP* numbers are summed up; the procedure is for calculating *FN*.

For Task 2, the labeled dataset includes disease images for the four pathogens, split into two datasets: 1324 images with masked ROIs and 1324 images with masked non-ROIs. For Task 3, the dataset includes 11 classes: four classes (3034 images) costing of grain-free images from the same four pathogens and seven classes (7000 images) consisting of images from seven healthy skin tissue types: subcutis, nerves, dermis, vessels, epidermis, muscle skeletal, and sebaceous. For both tasks, however, the split protocols and the evaluation measures are the same as in Task 1.

For the semantic image segmentation task (Task 4), the accuracy of the models is calculated using the Dice Coefficient and Intersection over Union (IoU) measures. The Dice Coefficient is a metric for evaluating the similarity between two sets, commonly used in semantic segmentation tasks. It measures the overlap between predicted and the expert-generated ground truth masks (2×1,324 masks in total, for both experts), ranging from 0 (no overlap) to 1 (perfect overlap). The Intersection over Union (IoU) measures the ratio of the intersection area to the union area of the predicted and ground truth masks.

The measures are calculated as:

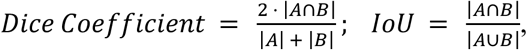

where A is the predicted segmentation mask (measured in pixels), and B is the ground truth segmentation mask. We note that two expert sources for ground truths are used (Mask1 and Mask2). Therefore, the accuracy measures are calculated for each ground truth independently.

For the last task of Explainable AI (Task 5), the performance can be assessed only for the images with detectable ROIs. Because no ground truth segmentation is available for the grain-free images, we will apply the same protocol and segmentation model that was assessed for the images with detectable ROIs, supported by the evaluation of the latter task. The evaluation of the accuracy of the saliency classification masks is carried out similarly to the evaluation of the segmentation task (Task 4). Specifically, we use the same test of 2×1,324 expert-generated ground truth masks and calculate the same *Dice Coefficient* and *IoU* scores.

The experimental framework was implemented using PyTorch (version 2.6.0+cu124) with all training and evaluation conducted on Tesla T4 GPUs through Google Colab’s cloud infrastructure. The computational environment leveraged mixed-precision training and gradient accumulation to optimize memory usage on the T4 GPAs (16GB VRAM). Training typically requires 6-12 hours per model configuration.

## Acknowledgments

This work is supported by the Regional Scholarship and Innovation Fund (RSIF) and the Partnerships for Skills in Applied Sciences, Engineering, and Technologies (PASET) to K.M.S.Z. The authors express their sincere gratitude to Dr. Emmanuel Edwar Siddig for his invaluable contributions to the annotation of histopathology slide samples.

