## Supplementary Materials for "Robust and accurate diagnosis of infectious skin diseases from histopathology images by integrating deep learning and explainable AI"

SUPPEMENTARY FIGURES

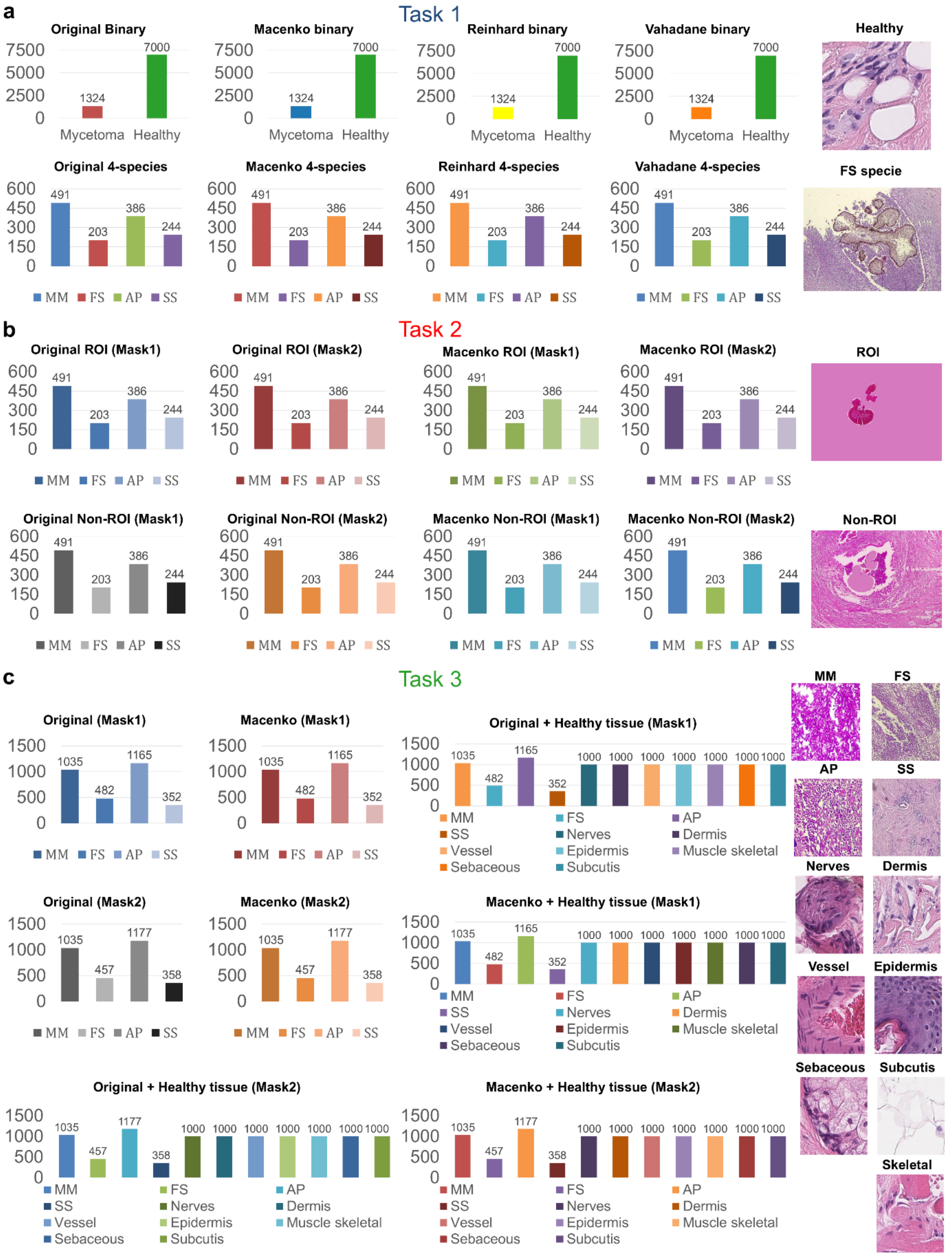


**Supplementary Figure S1. Dataset Composition Overview for Supervised Learning Tasks. (a) Task 1 Datasets: Binary and Multi-Class Classification**. This panel illustrates the eight datasets for Task 1. For binary classification (disease vs. healthy tissue), datasets combine 1,324 disease images with 7,000 healthy tissues. For multi-class classification of four pathogenic species, only the 1,324 disease images are used. All datasets include raw and normalized (Macenko, Reinhard, Vahadane) image versions. **(b) Task 2 Datasets: ROI vs. Non-ROI Impact.** This panel shows the eight paired datasets for Task 2, comprising 1,324 Region of Interest (ROI) images and 1,324 corresponding Non-ROI images. These datasets were derived from both original and Macenko-normalized images, utilizing Mask1 and Mask2 annotations. **(c) Task 3 Datasets: Grain-Free Image Classification.** This panel presents the eight datasets for Task 3, designed for grain-free image classification. Datasets contain 3,034 (Mask1) or 3,027 (Mask2) grain-free fragments from raw and Macenko-normalized specimens. Some datasets also incorporate 7,000 healthy tissue images as negative controls.

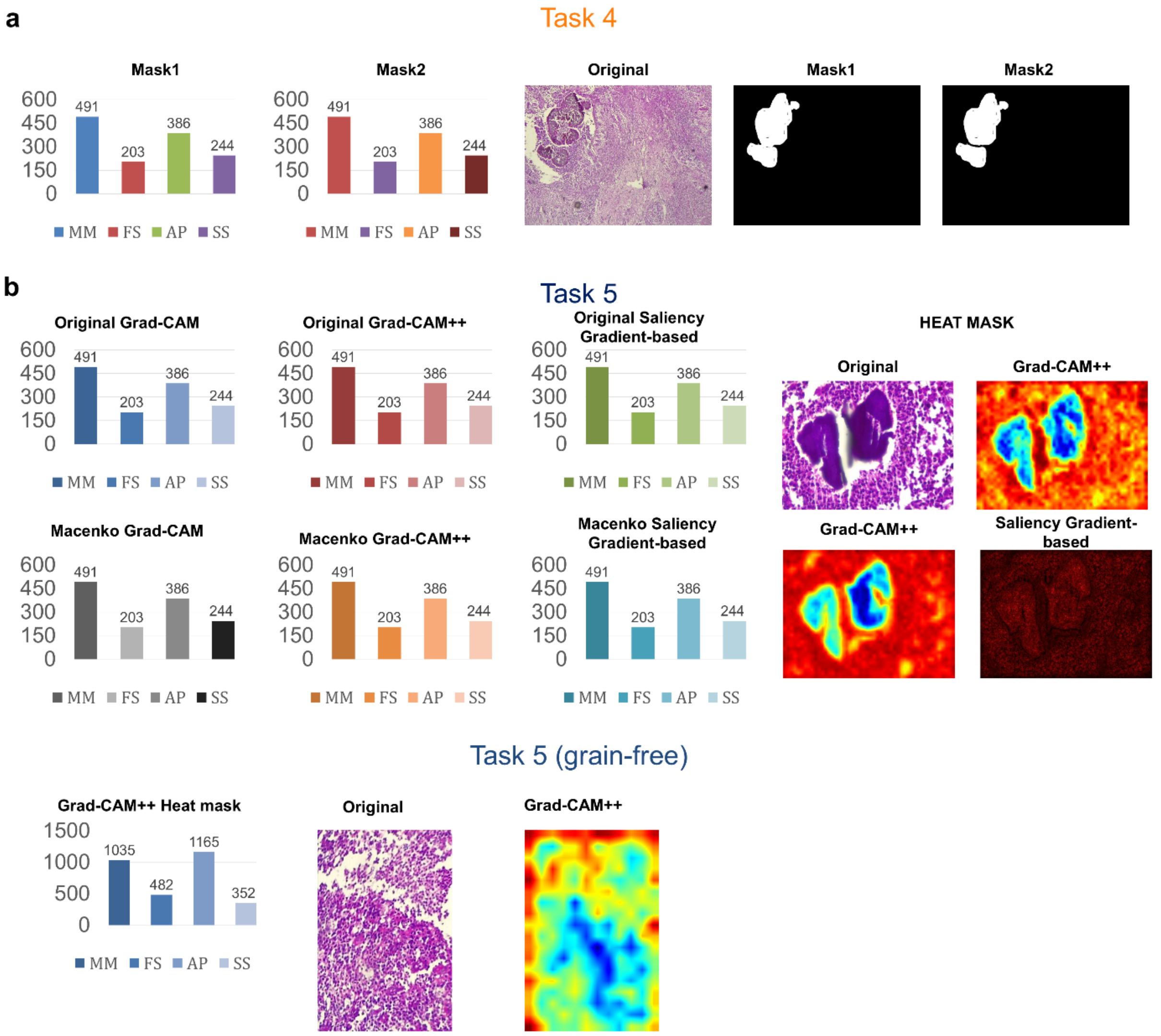


**Supplementary Figure S2: Dataset Composition for Semantic Segmentation and Explainable AI. (a) Task 4 Datasets: Semantic Image Segmentation Ground Truths**. This panel illustrates the two primary ground truth datasets for Task 4. These datasets consist of manual annotations (Mask1 and Mask2), serving as distinct ground truth sets for training and evaluating semantic segmentation models. **(b) Task 5 Datasets: Explainable AI Heat Mask Generation.** This panel details the seven datasets of heat masks generated for Task 5. Six datasets comprise heat masks derived from raw and Macenko-normalized images using GradCAM, GradCAM++, and Saliency gradient-based techniques. The seventh dataset specifically includes heat masks for grain-free images, generated using GradCAM++.

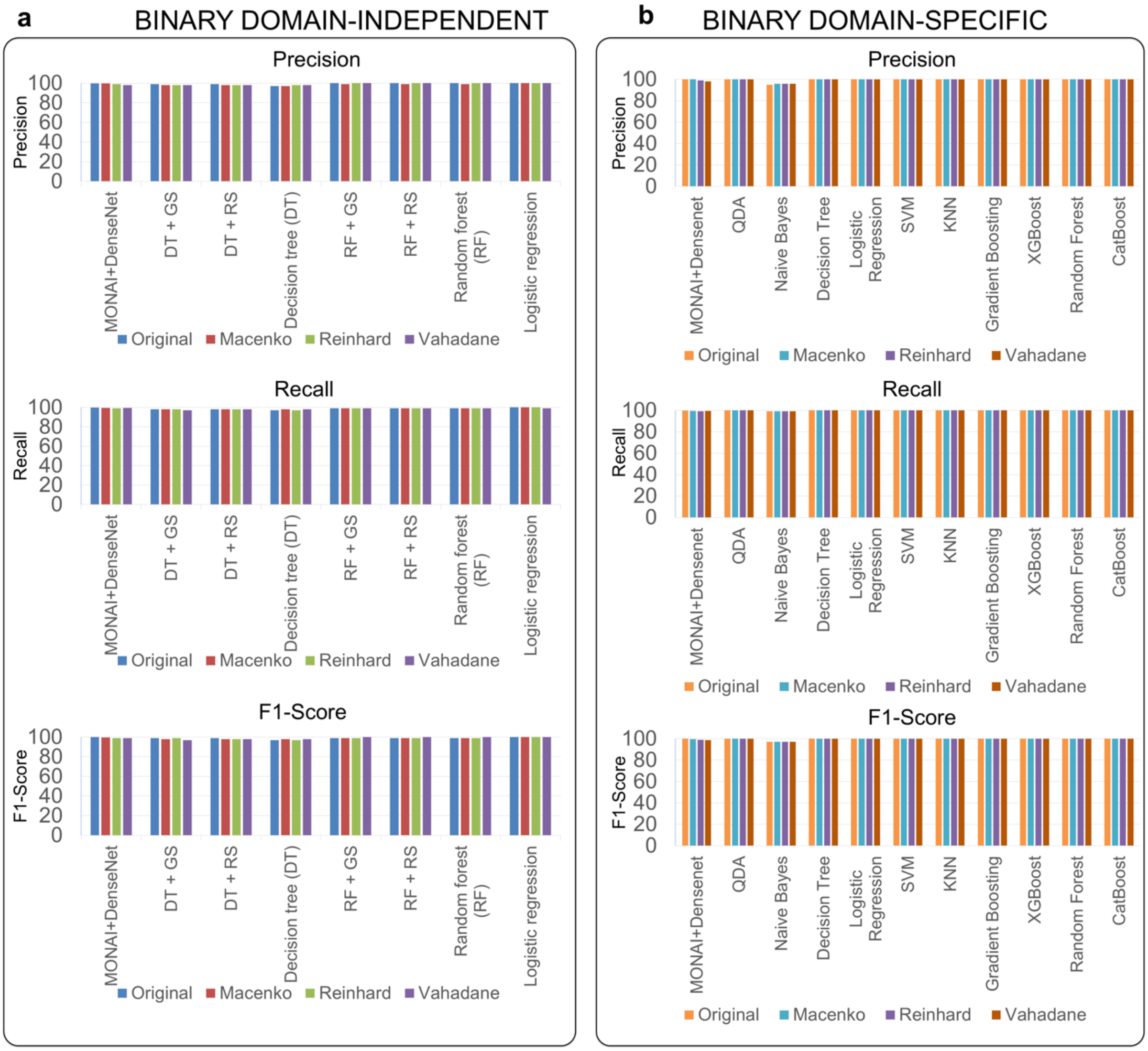


**Supplementary Figure S3: Performance Metrics for Binary Classification Tasks. (a) Binary Domain-Independent Metric Plot.** This panel displays the precision, recall, and F1-score for a range of linear, non-linear, and deep learning models. Logistic Regression consistently achieved the highest scores, often reaching 100% precision, recall, and F1-score across all normalization methods. MONAI+DenseNet also demonstrated excellent performance, particularly on original and Macenko-normalized images. **(b) Binary Domain-Specific Metric Plot.** This panel shows the precision, recall, and F1-score for models optimized for domain-specific characteristics. Most models in this category, including QDA, Decision Tree, Logistic Regression, SVM, KNN, Gradient Boosting, XGBoost, Random Forest, and CatBoost, achieved perfect (100%) precision, recall, and F1-scores for many normalized datasets. Naive Bayes showed slightly lower, but still strong, performance. These results highlight the high discriminative power of these models for domain-specific binary classification when appropriate normalization is applied.

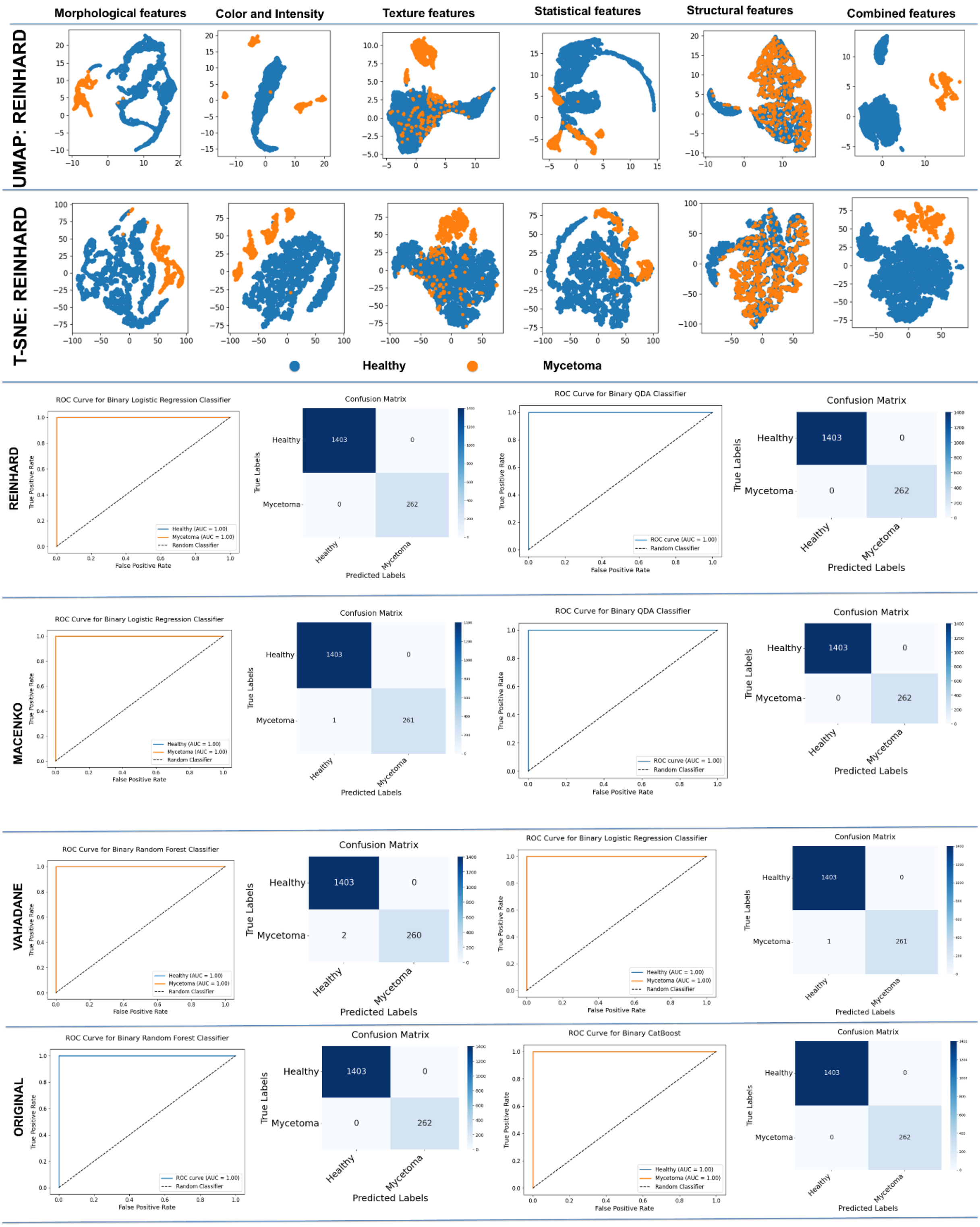


**Supplementary Figure S4: Domain-Specific Feature Visualization and Shallow Classifier Performance for Binary Classification. (a) UMAP Visualization.** This panel displays UMAP plots illustrating class separability based on individual and combined domain-specific features (morphological, color/intensity, texture, structural) using the Reinhard-normalized dataset. **(b) t-SNE Visualization.** This panel presents t-SNE plots, offering an alternative dimensionality reduction view on the same domain-specific features and their ability to discriminate classes with the Reinhard-normalized dataset. **(c) Reinhard ROC Curve and Confusion Matrix.** This panel shows the ROC curve and confusion matrix for Logistic Regression and QDA shallow classifiers, demonstrating their binary classification performance on the Reinhard-normalized dataset. **(d) Macenko ROC Curve and Confusion Matrix.** This panel presents the ROC curve and confusion matrix for Logistic Regression and QDA shallow classifiers, evaluating their binary classification performance on the Macenko-normalized dataset. **(e) Vahadane ROC Curve and Confusion Matrix.** This panel displays the ROC curve and confusion matrix for Random Forest and Logistic Regression shallow classifiers, illustrating their binary classification performance on the Vahadane-normalized dataset. **(f) Original ROC Curve and Confusion Matrix.** This panel shows the ROC curve and confusion matrix for Random Forest and CatBoost shallow classifiers, evaluating their binary classification performance on the original (raw) dataset.

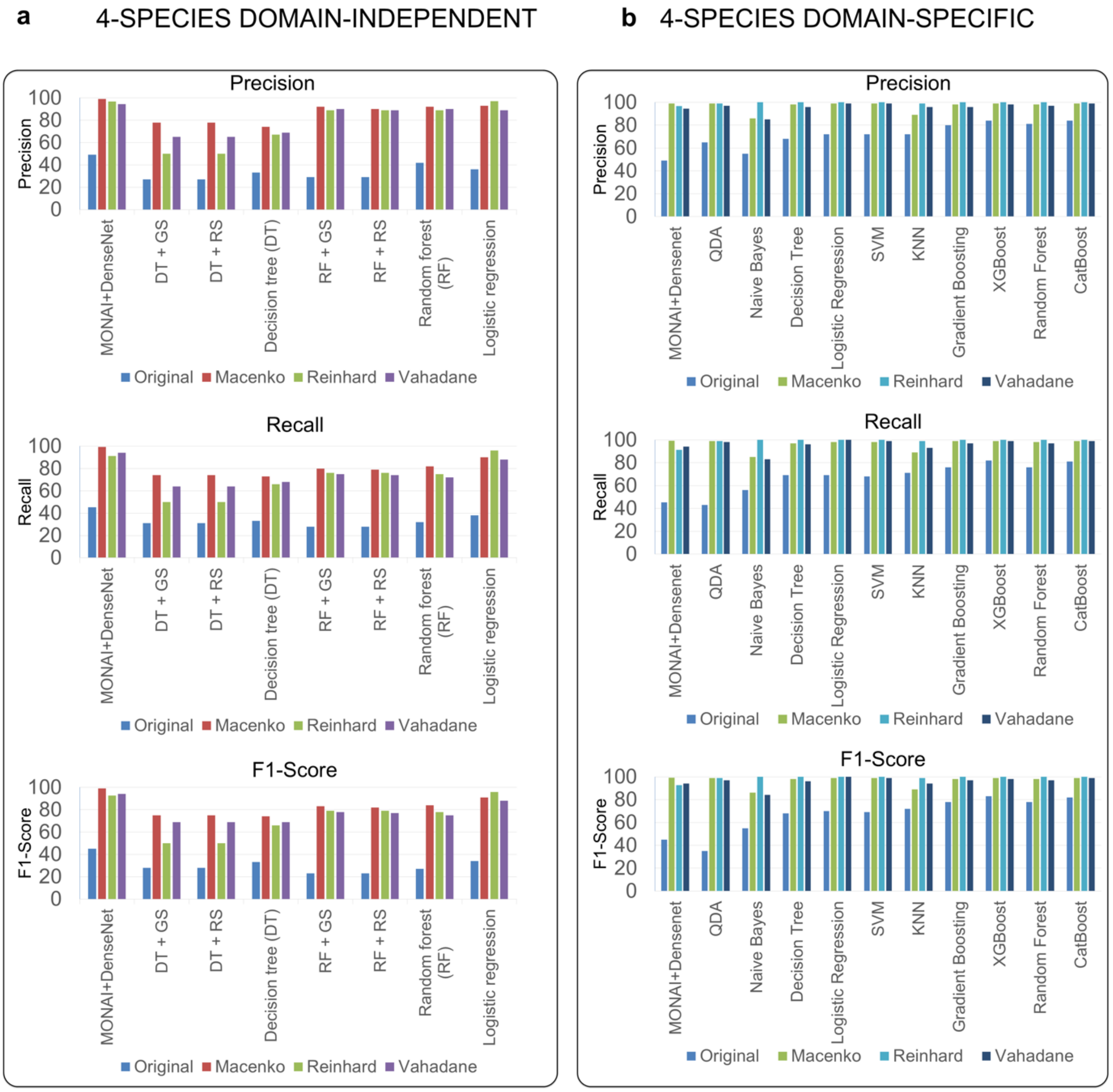


**Supplementary Figure S5: Performance Metrics for 4-Species Multi-Class Classification Tasks. (a) 4-Species Domain-Independent Metric Plot.** This panel displays the precision, recall, and F1-score for a range of linear, non-linear, and deep learning models. MONAI+DenseNet consistently demonstrated superior performance across all metrics and normalization methods, achieving remarkably high F1-scores, particularly with Macenko-normalized images (99.20%). Among the shallow learning models, Logistic Regression, RF + GS, and Random Forest also showed strong F1-scores, frequently reaching 89-97% when combined with Macenko, Reinhard, or Vahadane normalization, highlighting the significant benefits of normalization for multi-class classification. Performance on original (raw) images was notably lower for most models compared to normalized data, emphasizing the importance of preprocessing. **(b) 4-Species Domain-Specific Metric Plot.** This panel shows the precision, recall, and F1-score for models optimized for domain-specific characteristics. A substantial number of these models, including Logistic Regression, SVM, KNN, Gradient Boosting, XGBoost, Random Forest, and CatBoost, achieved exceptional F1-scores, frequently reaching 98%, 99%, or even 100% when applied to Macenko-normalized, Reinhard-normalized, or Vahadane-normalized images. QDA also showed very strong performance with normalized data. This consistently high performance underscores the robust capabilities of these models for distinguishing between the four pathogenic species, especially when combined with effective image normalization techniques.

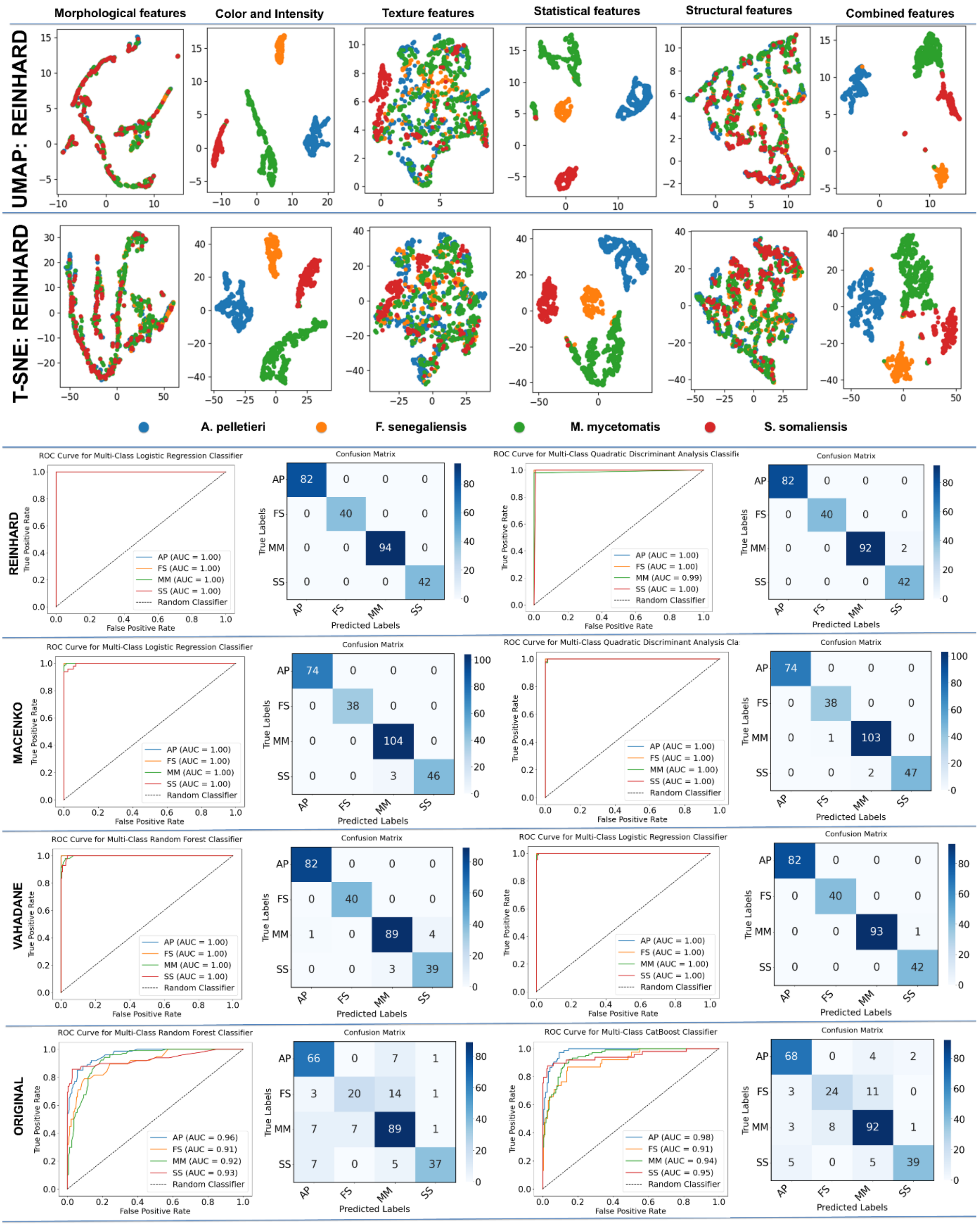


**Supplementary Figure S6: Domain-Specific Feature Visualization and Shallow Classifier Performance for 4-Species Classification. (a) UMAP Visualization.** This panel displays UMAP plots illustrating the separability of the four species based on individual and combined domain-specific features (morphological, color/intensity, texture, structural) using the Reinhard-normalized dataset. **(b) t-SNE Visualization.** This panel presents t-SNE plots, offering an alternative dimensionality reduction view on the same domain-specific features and their ability to discriminate species with the Reinhard-normalized dataset. **(c) Reinhard ROC Curve and Confusion Matrix.** This panel shows the ROC curve and confusion matrix for Logistic Regression and QDA shallow classifiers, demonstrating their 4-species classification performance on the Reinhard-normalized dataset. **(d) Macenko ROC Curve and Confusion Matrix.** This panel presents the ROC curve and confusion matrix for Logistic Regression and QDA shallow classifiers, evaluating their 4-species classification performance on the Macenko-normalized dataset. **(e) Vahadane ROC Curve and Confusion Matrix.** This panel displays the ROC curve and confusion matrix for Random Forest and Logistic Regression shallow classifiers, illustrating their 4-species classification performance on the Vahadane-normalized dataset. **(f) Original ROC Curve and Confusion Matrix.** This panel shows the ROC curve and confusion matrix for Random Forest and CatBoost shallow classifiers, evaluating their 4-species classification performance on the original (raw) dataset.

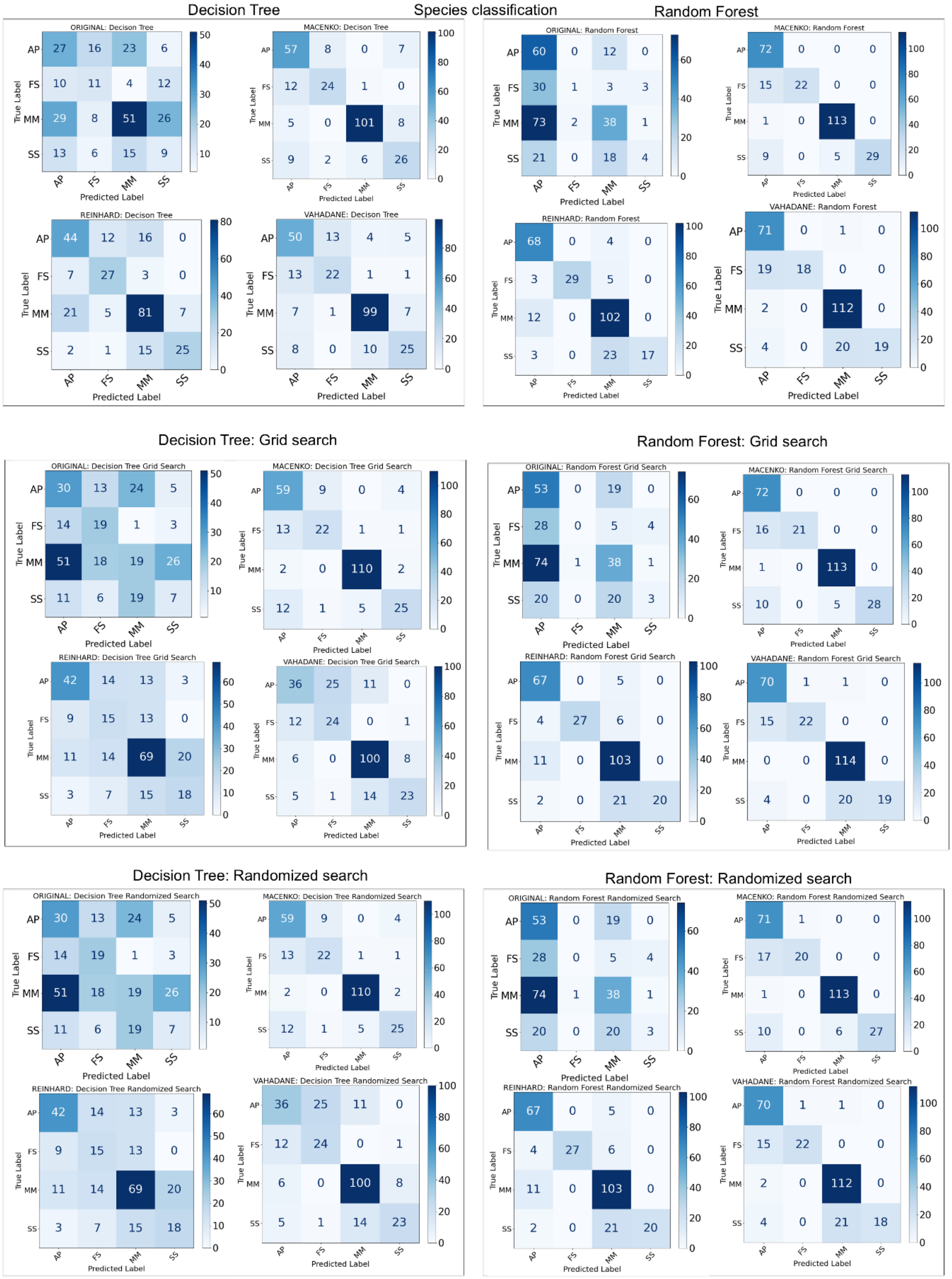


**Supplementary Figure S7: Confusion Matrices for 4-Species Classification with Domain-Independent Features.** This figure presents the confusion matrices detailing the performance of various classifiers on the 4-species classification task using domain-independent features. Each matrix illustrates the true versus predicted labels for different combinations of models and image normalization methods. Results are shown for Decision Tree (simple, Grid Search optimized, Randomized Search optimized) and Random Forest (simple, Grid Search optimized, Randomized Search optimized) applied to raw,

Macenko-normalized, Reinhard-normalized, and Vahadane-normalized images. These matrices visually represent the accuracy and types of misclassifications for each classifier and normalization strategy

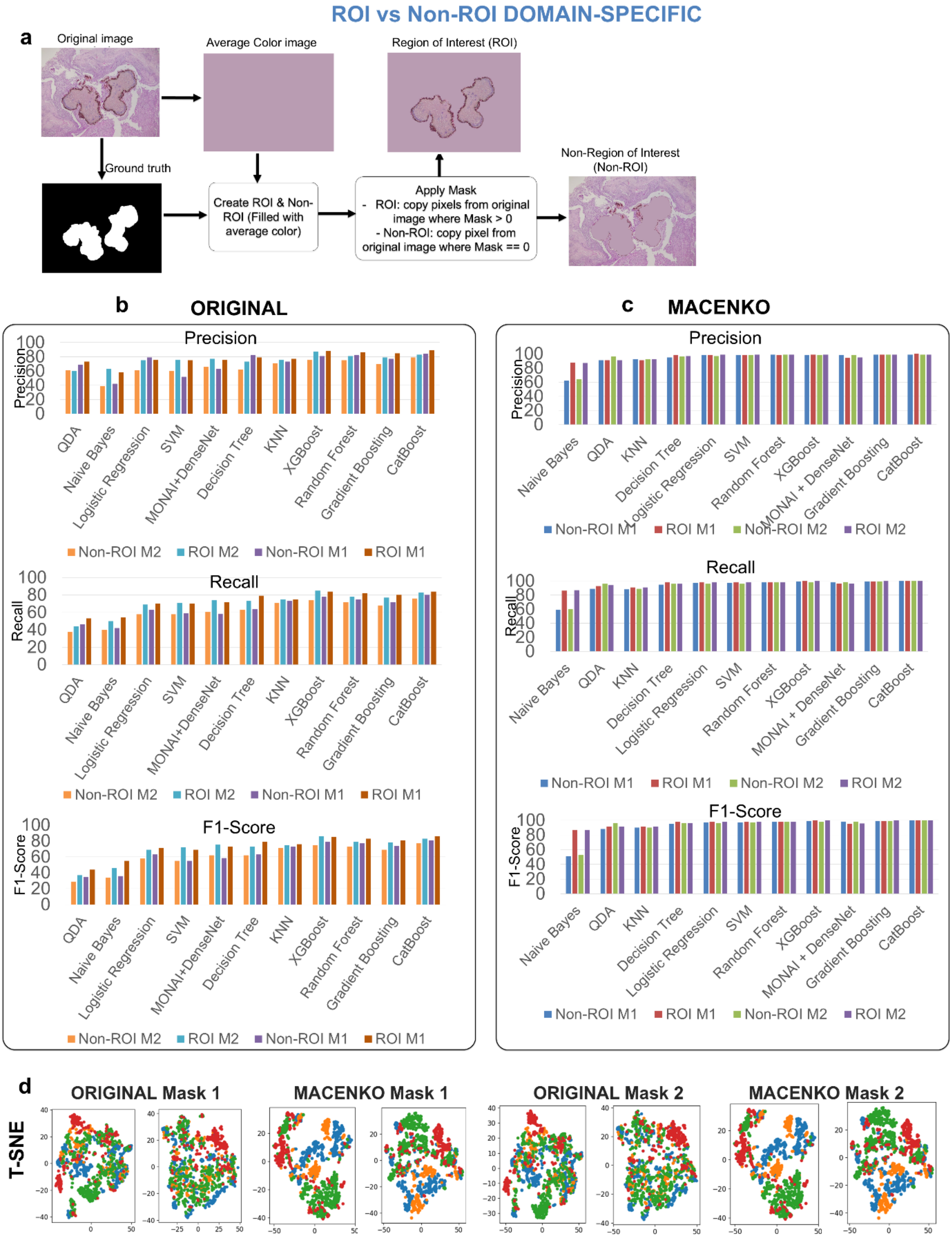


**Supplementary Figure S8: Impact of ROI and Normalization on 4-Species Classification Performance and Feature Visualization**. **(a) ROI and Non-ROI Dataset Generation Workflow.** This panel illustrates the process for creating ROI-isolated and Non-ROI datasets using Mask1 and Mask2 annotations. **(b) Performance Metrics: Original Images.** This panel presents precision, recall, and F1-scores for models on original ROI and Non-ROI images. ROI datasets generally yield higher F1-scores, with XGBoost, Random Forest, and CatBoost showing top performance. **(c)**

**Performance Metrics: Macenko-Normalized Images.** This panel displays metrics for models on Macenko-normalized images. A dramatic performance improvement is observed across nearly all models with normalization, with numerous models achieving 98-100% F1-scores. CatBoost, XGBoost, and Gradient Boosting consistently show near-perfect scores across all conditions. **(d) t-SNE Visualization of Four-Species Grouping.** This panel presents t-SNE plots visualizing learned features for the four species. These plots illustrate species separability with original versus Macenko-normalized images and the influence of Mask1/Mask2 annotations.

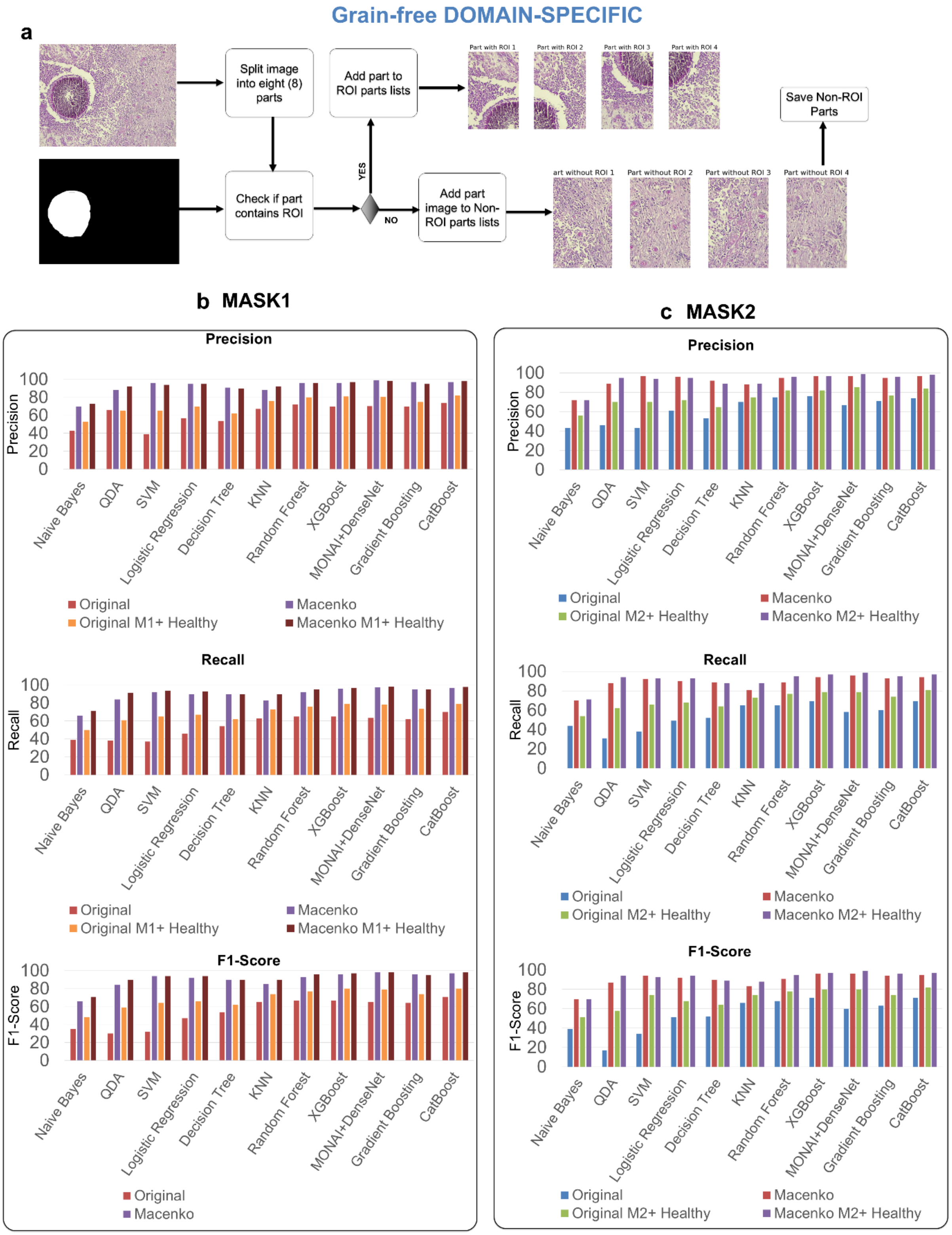


**Supplementary Figure S9: Grain-Free Dataset Generation and Classification Performance**. **(a) Generation of Grain-Free Datasets Workflow.** This panel outlines the methodology used to create the grain-free image datasets. It should visually explain how image fragments without visible grains were extracted using Mask1 and Mask2 annotations. **(b) Mask 1: Grain-Free Classification Performance.** This panel presents precision, recall, and F1-scores for models on grain-free datasets derived using Mask1, comparing raw, Macenko-normalized, and their combinations with healthy tissue images. Macenko normalization consistently yields substantial performance improvements (e.g., F1-scores for MONAI+DenseNet jumping from 65.24% to 98.33% on grain-free images). The inclusion of healthy tissue images further enhances F1-scores, particularly for original (raw) data. MONAI+DenseNet, XGBoost, and CatBoost consistently show the highest F1-scores (e.g., 97-98% for Macenko-normalized data with healthy controls), demonstrating robust performance on these challenging samples. **(c) Mask 2: Grain-Free Classification Performance.** This panel displays similar performance metrics for grain-free datasets generated with Mask2. Similar to Mask1, Macenko normalization dramatically improves performance across all models and the inclusion of healthy controls further boosts scores. MONAI+DenseNet consistently achieves the top F1-score (99% for Macenko M2 + Healthy), while XGBoost, Random Forest, Gradient Boosting, and CatBoost also show excellent performance (95-97% F1-scores) on normalized data with healthy controls. This reinforces the importance of both normalization and negative controls for accurate grain-free classification.

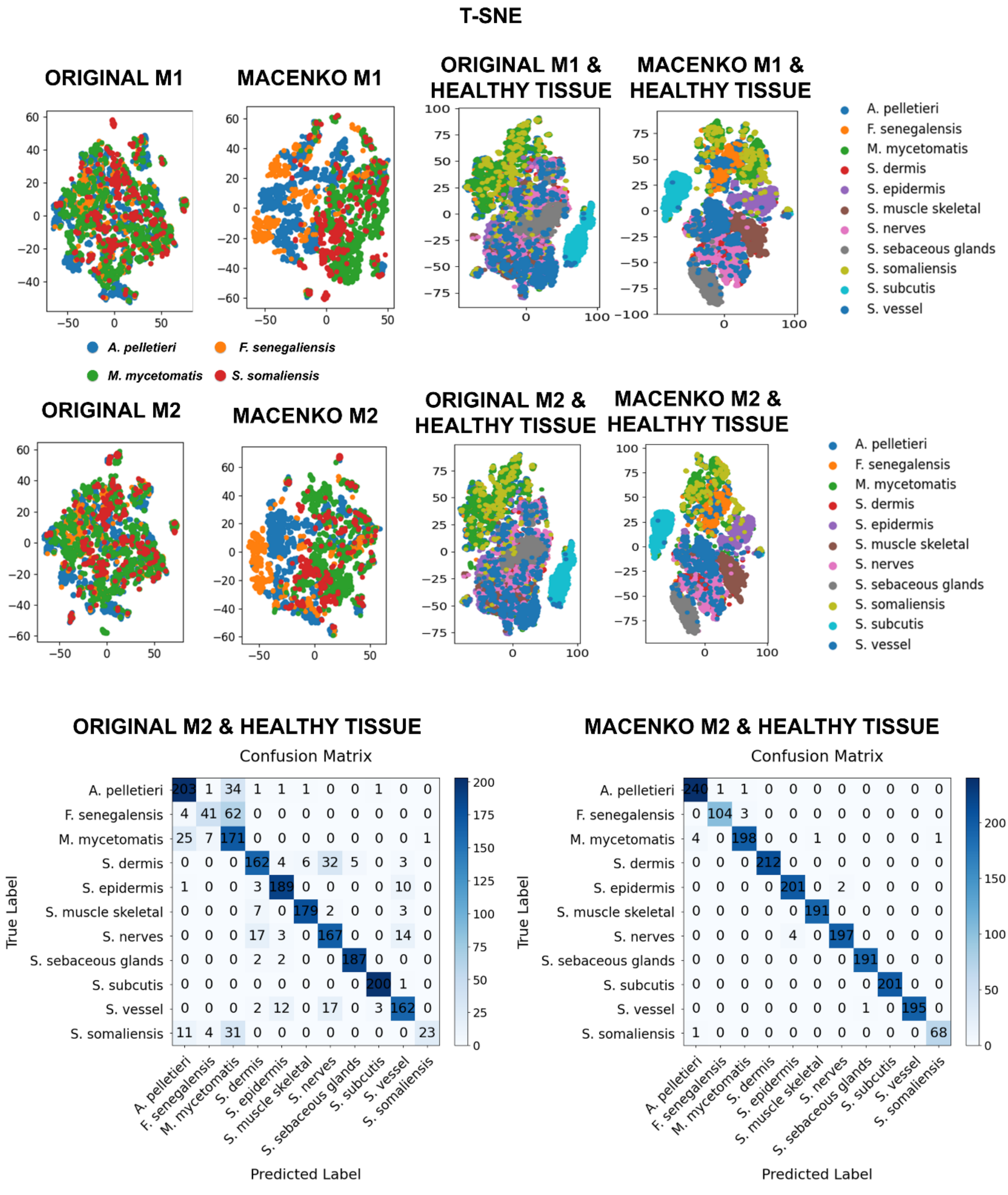


**Supplementary Figure S10: Grain-Free Feature Visualization and Classification Performance. (a) t-SNE Visualization with Mask 1.** This panel displays t-SNE plots for Mask 1-segmented grain-free images (raw, Macenko-normalized, and with healthy tissue). Raw images show poor separation; Macenko normalization offers minor improvements. Significant class separability emerges when Macenko-normalized images are combined with healthy tissue, reflecting improved classification. **(b) t-SNE Visualization with Mask 2.** This panel presents t-SNE plots for Mask 2-segmented grain-free images under similar conditions. Similar to Mask 1, optimal class distinction is achieved when Macenko-normalized images are combined with healthy tissue controls, highlighting the dual benefits of preprocessing and negative controls. **(c) Confusion Matrices for Mask 2 with Healthy Tissue for MONAI-Densenet.** This panel provides confusion matrices comparing raw and Macenko-normalized

Mask 2 images, both combined with healthy tissue. These matrices quantitatively confirm that Macenko normalization drastically improves classification accuracy when healthy tissue controls are included.

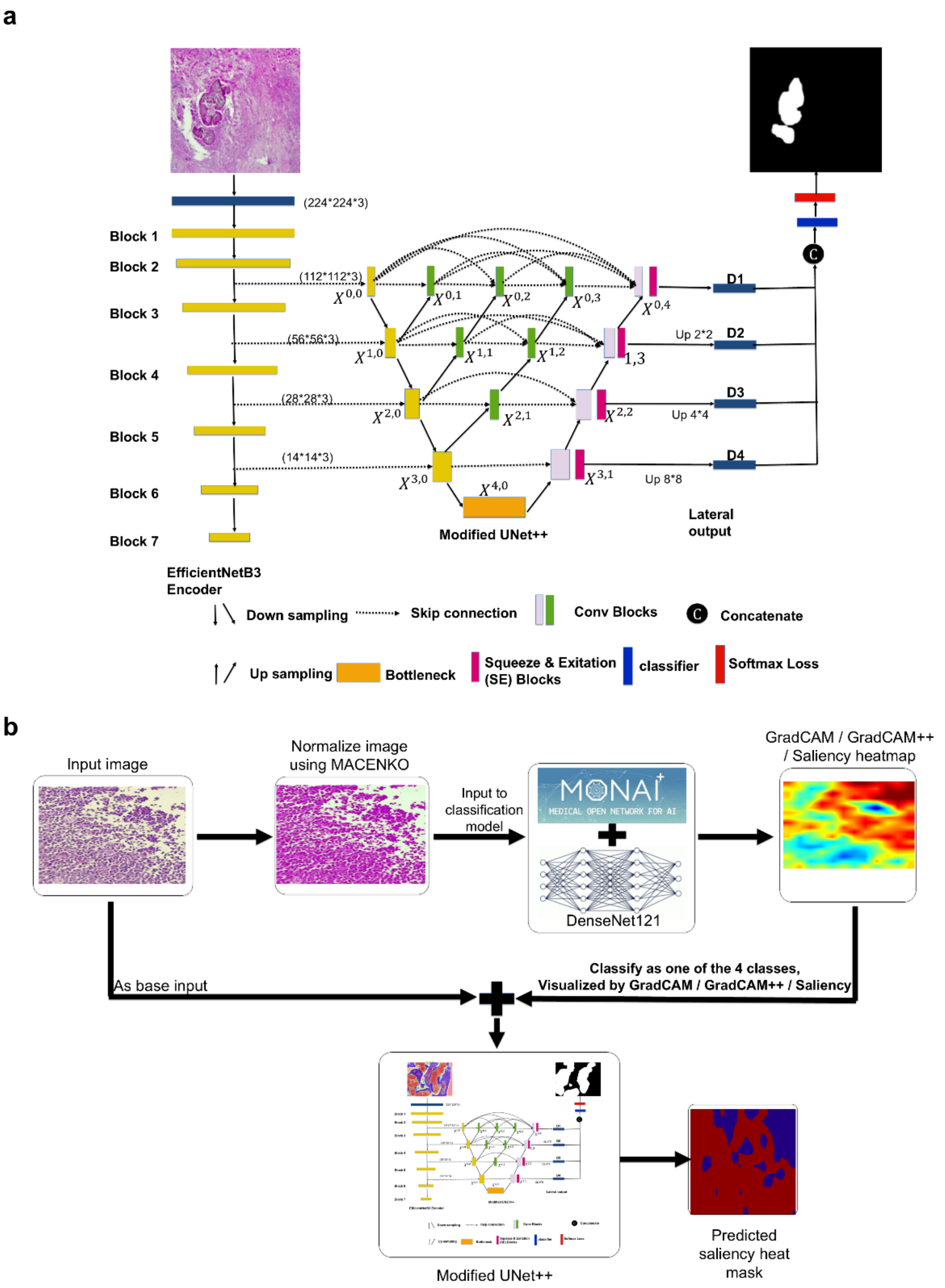


**Supplementary Figure S11: Workflows for Semantic Segmentation and Explainable AI Tasks. (a) Semantic Segmentation Workflow.** This panel details the process of performing semantic segmentation using the modified UNet++ architecture. It highlights the integration of an EfficientNet-

B3 backbone for feature extraction and the application of Spatial and Channel Squeeze & Excitation (SCSE) modules for optimized performance. **(b) Explainable AI Task Workflow.** This panel outlines the workflow for conducting the explainable AI task for grain-free images. It visually describes the steps involved in generating and processing saliency maps to provide insights into model decision-making, from input image to interpretable output.

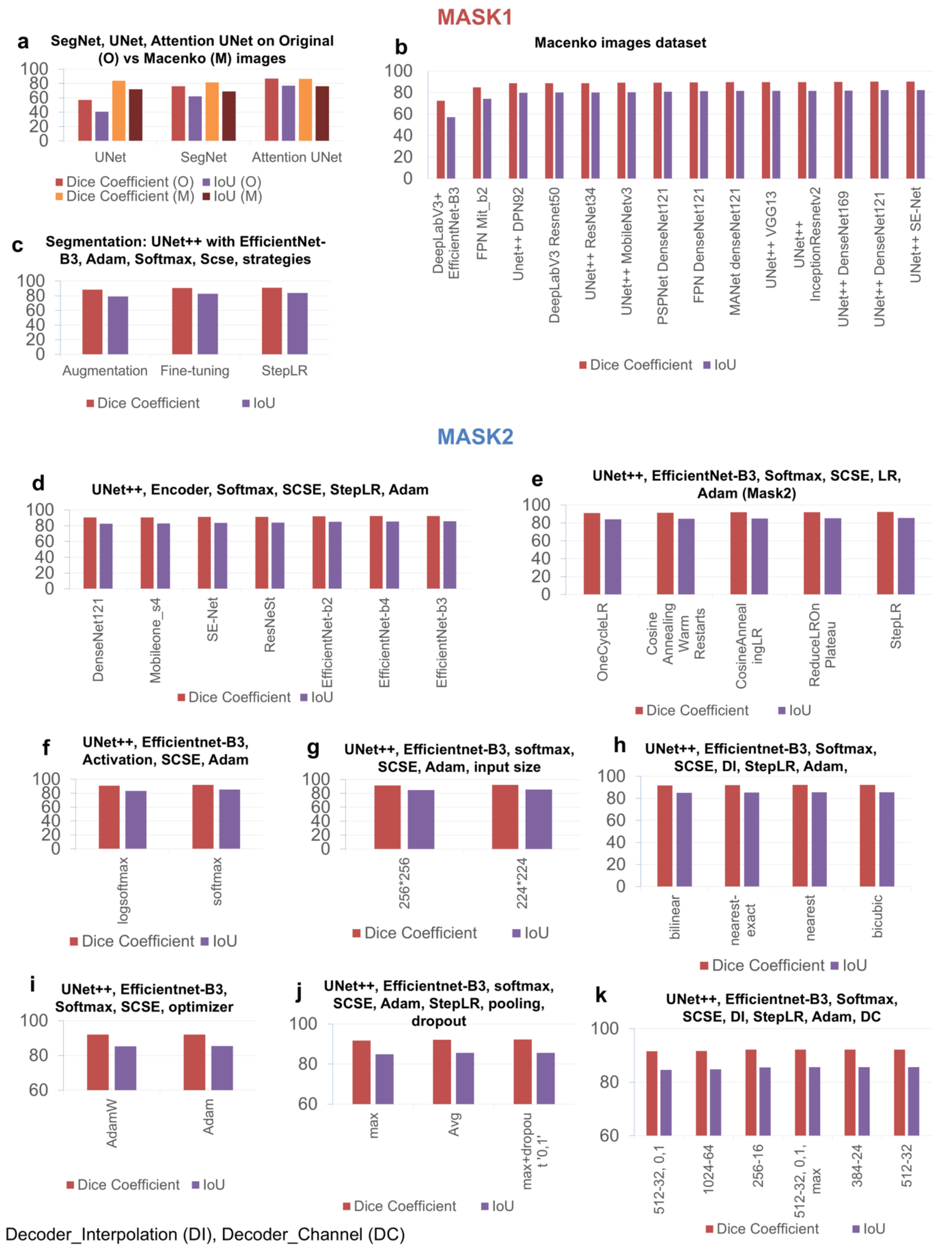


**Supplementary Figure S12: Comprehensive Semantic Segmentation Model Performance Evaluation. Mask 1 Performance (a)-(c). (a) Basic Model Comparison (Original vs. Macenko).** This panel compares the Dice Coefficient and IoU of UNet, SegNet, and Attention UNet on original vs. Macenko-normalized images. Attention UNet consistently shows superior performance on original images, while all models show significant improvement with Macenko normalization. **(b) Advanced Architecture Comparison (Macenko).** This panel evaluates a range of advanced segmentation models on Macenko-normalized images. UNet++ architectures, particularly with Densenet121, SE-Net, and Efficientnet-b3 backbones, consistently achieve the highest Dice Coefficients (e.g., UNet++Efficientnet-b3 at 90.54%). **(c) Training Strategy Impact.** This panel compares the effect of augmentation, fine-tuning, and StepLR on UNet++ with EfficientNet-B3. StepLR yields the highest Dice Coefficient (91.13%), indicating its effectiveness. **Mask 2 Performance for UNet++ with EfficientNet-B3, Softmax, SCSE, and Adam, unless specified (d)-(k). (d) Encoder Backbones.** This panel compares different encoder backbones. Efficientnet-b3 and efficientnet-b4 achieve the highest Dice Coefficients (92.16% and 92.12% respectively), demonstrating their strong feature extraction capabilities. **(e) Learning Rate Schedulers.** This panel evaluates various learning rate schedulers. StepLR again shows the top performance (92.16% Dice Coefficient), confirming its robustness for this task. **(f) Activation Functions.** This panel compares logsoftmax and softmax activations. Softmax significantly outperforms logsoftmax (92.08% Dice Coefficient). **(g) Input Size.** This panel assesses the impact of input image resolution. The 224x224 input size achieves a slightly higher Dice Coefficient (92.08%) than 256x256. **(h) Decoder Interpolation (DI).** This panel compares different interpolation methods used in the decoder. Bicubic interpolation yields the highest Dice Coefficient (92.19%). **(i) Optimizers.** This panel presents results for AdamW. AdamW shows a strong Dice Coefficient of 92.02%. **(j) Pooling and Dropout.** This panel explores pooling strategies and dropout. Average pooling combined with dropout (0.1) provides the highest Dice Coefficient (92.21%). **(k) Decoder Channel (DC) and Deep Supervision.** This panel shows the impact of various decoder channel configurations and deep supervision. The 384-24 configuration, along with the 512-32 (max) and 512-32 (no max) setups, demonstrate the highest Dice Coefficients (92.21%, 92.20%, 92.23% respectively)**,** indicating the benefits of deep supervision for robust segmentation.

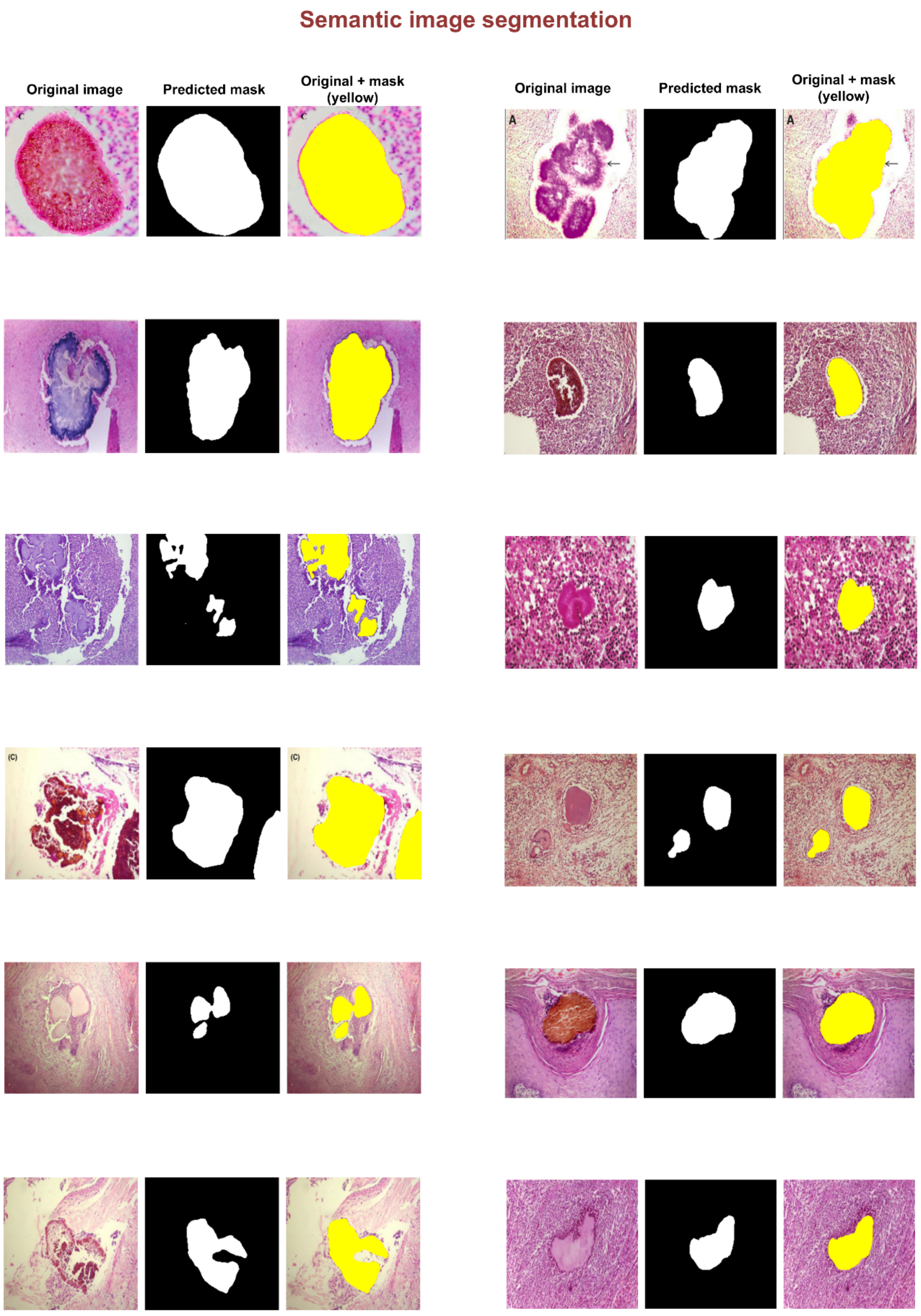


**Supplementary Figure S13: Automatic Semantic Image Segmentation Results (Examples 1-12).** This figure presents examples of automatic semantic image segmentation, demonstrating the framework's ability to delineate regions of interest without manual intervention. For each input image, the display includes the predicted mask, representing the automatically generated segmentation, followed by the overlay of the original image with the predicted mask. As these images lack pre-existing ground truth annotations, these visualizations highlight the model's independent segmentation capabilities on unseen data.

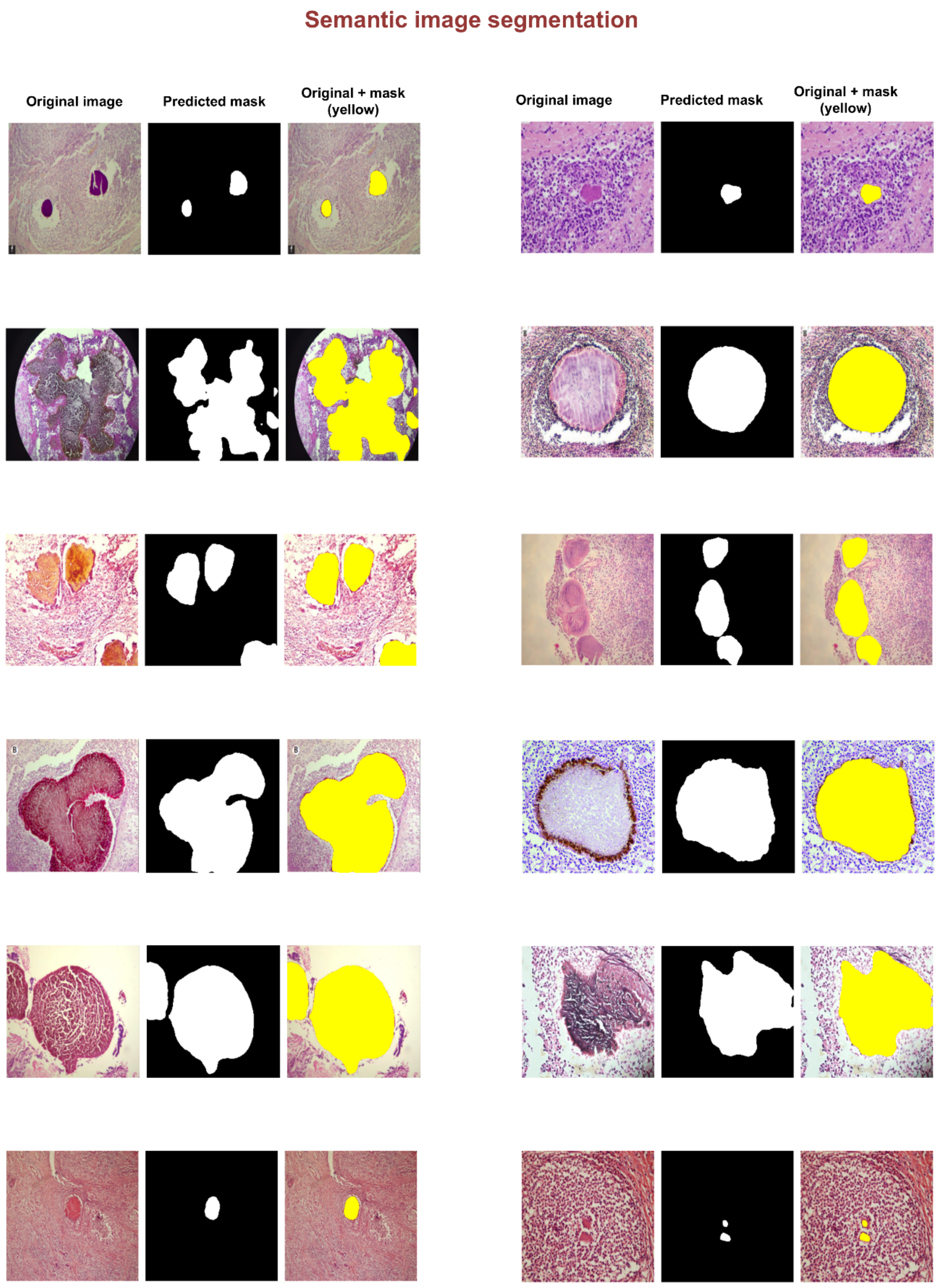


**Supplementary Figure S14: Automatic Semantic Image Segmentation Results (Examples 13-24).** This figure presents examples of automatic semantic image segmentation, demonstrating the framework's ability to delineate regions of interest without manual intervention. For each input image, the display includes the predicted mask, representing the automatically generated segmentation, followed by the overlay of the original image with the predicted mask. As these images lack pre-existing ground truth annotations, these visualizations highlight the model's independent segmentation capabilities on unseen data.

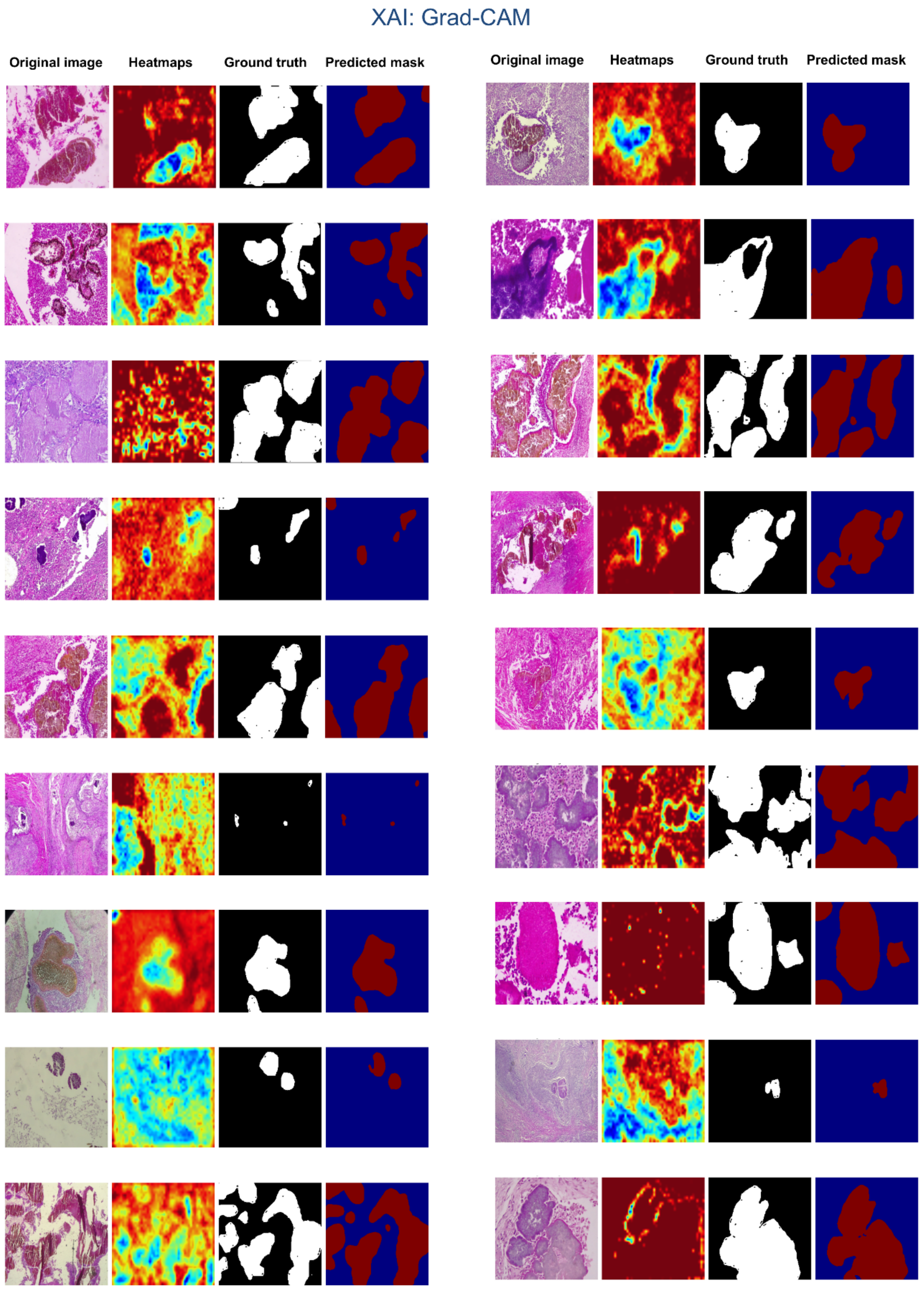


**Supp Figure 15. Supplementary Figure S15: Grad-CAM Analysis with Ground Truth for Explainable 4-Species Classification.** This figure presents representative examples of Grad-CAM analysis on original 4-species histopathology images, serving as an Explainable AI task to provide model interpretability. For each sample, the display includes the original 4-species image, its saliency heat mask (highlighting classifier-informative regions), the corresponding manually annotated ground truth mask (Mask1 or Mask2) from pathologists, and a binary saliency classification mask derived from the heat map. This visualization compares the model's focus with true pathological regions, demonstrating alignment between classifier decisions and expert annotations.

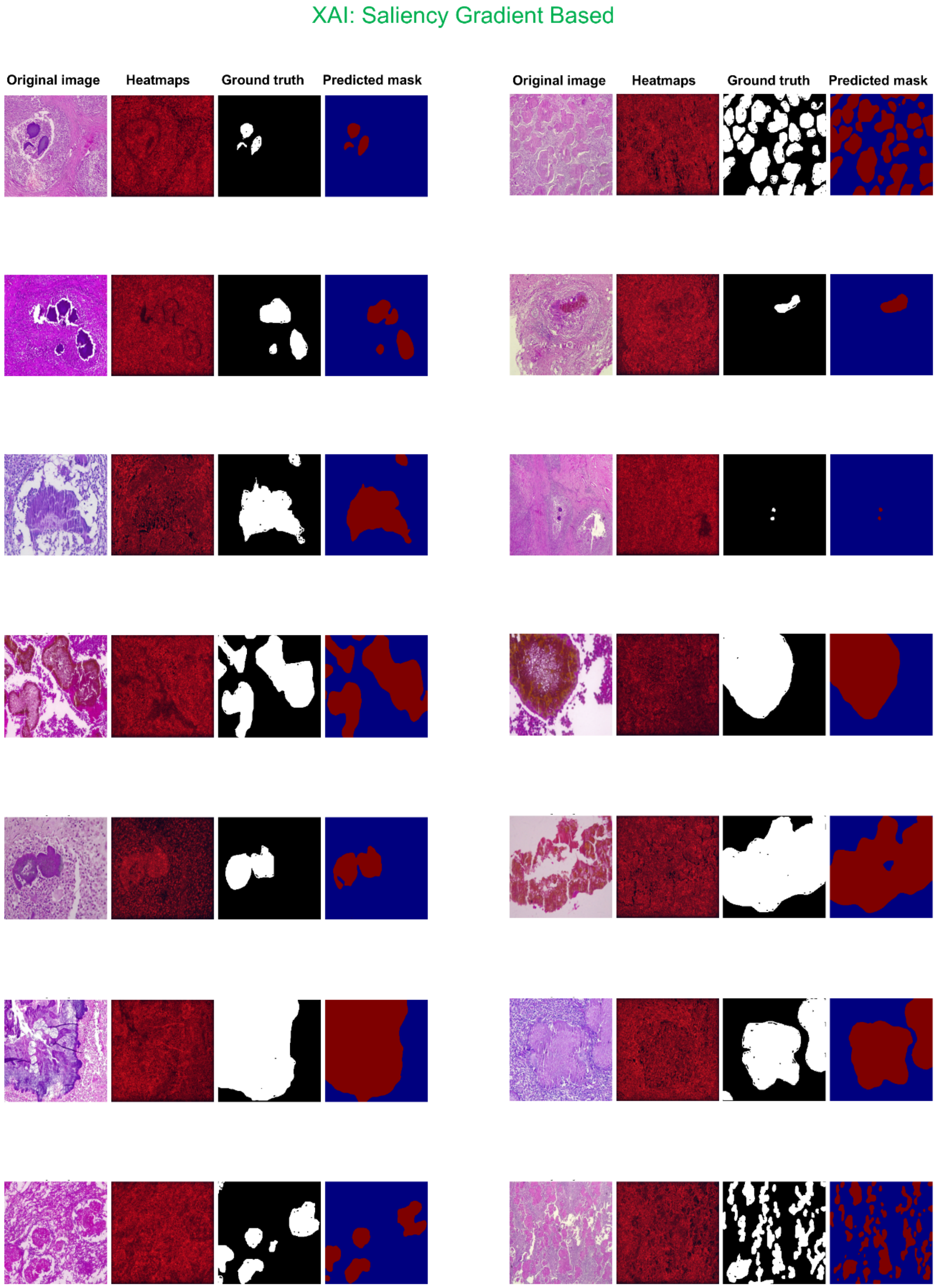


**Supplementary Figure S16: Saliency Gradient-Based Analysis with Ground Truth for Explainable 4-Species Classification**. This figure presents representative examples of saliency gradient-based analysis on original 4-species histopathology images, serving as an Explainable AI task to provide model interpretability. For each sample, the display includes the original 4-species image, its saliency heat mask (highlighting classifier-informative regions), the corresponding manually annotated ground truth mask (Mask1 or Mask2) from pathologists, and a binary saliency classification mask derived from the heat map. This visualization compares the model's focus with true pathological regions, demonstrating alignment between classifier decisions and expert annotations.

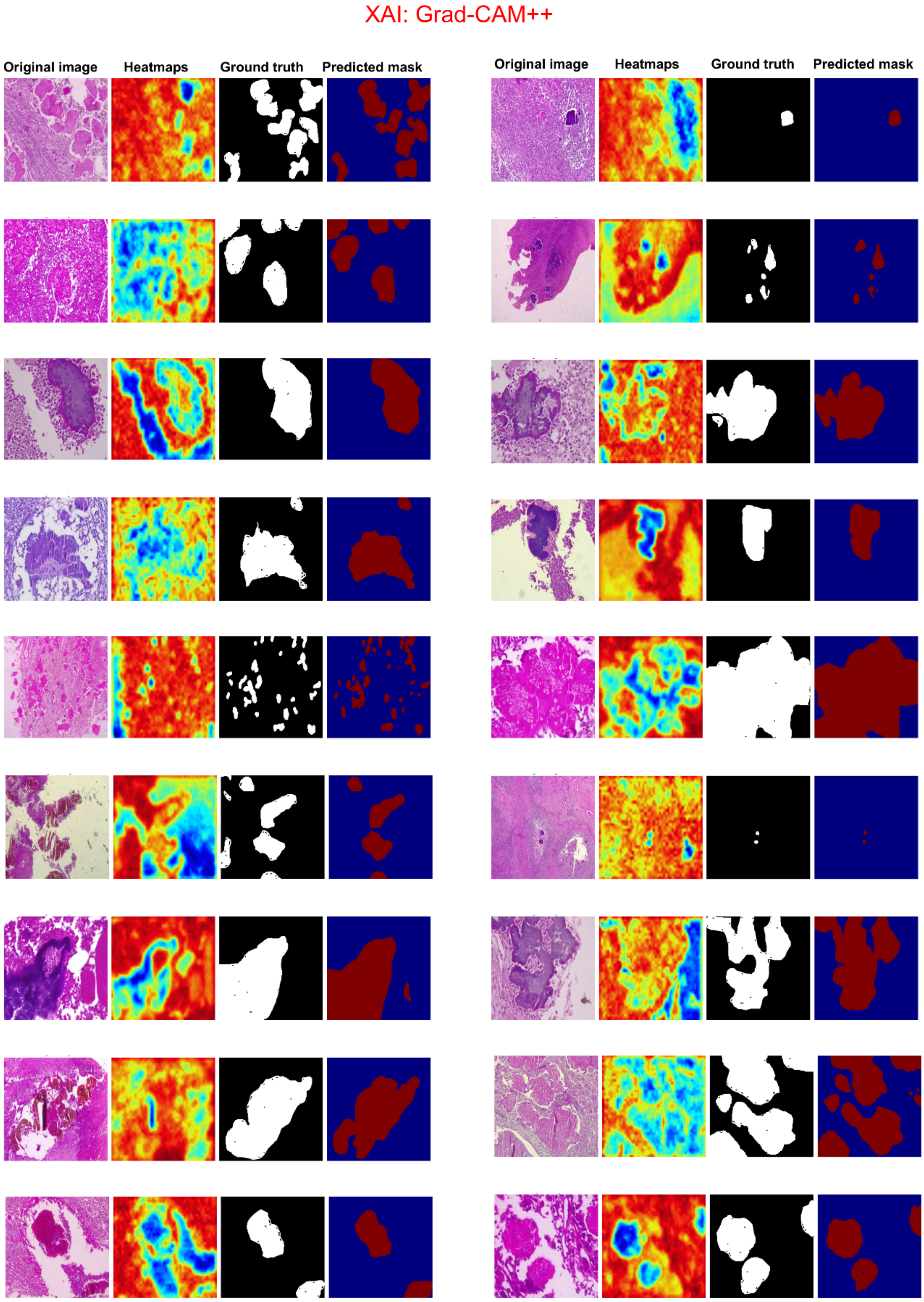


**Supplementary Figure S17: Grad-CAM++ Saliency Analysis with Ground Truth for Explainable 4-Species Classification.** This figure presents representative examples of Grad-CAM++ saliency analysis on original 4-species histopathology images, serving as an Explainable AI task to provide model interpretability. For each sample, the display includes the original 4-species image, its saliency heat mask (highlighting classifier-informative regions), the corresponding manually annotated ground truth mask (Mask1 or Mask2) from pathologists, and a binary saliency classification mask derived from the heat map. This visualization compares the model's focus with true pathological regions, demonstrating alignment between classifier decisions and expert annotations.

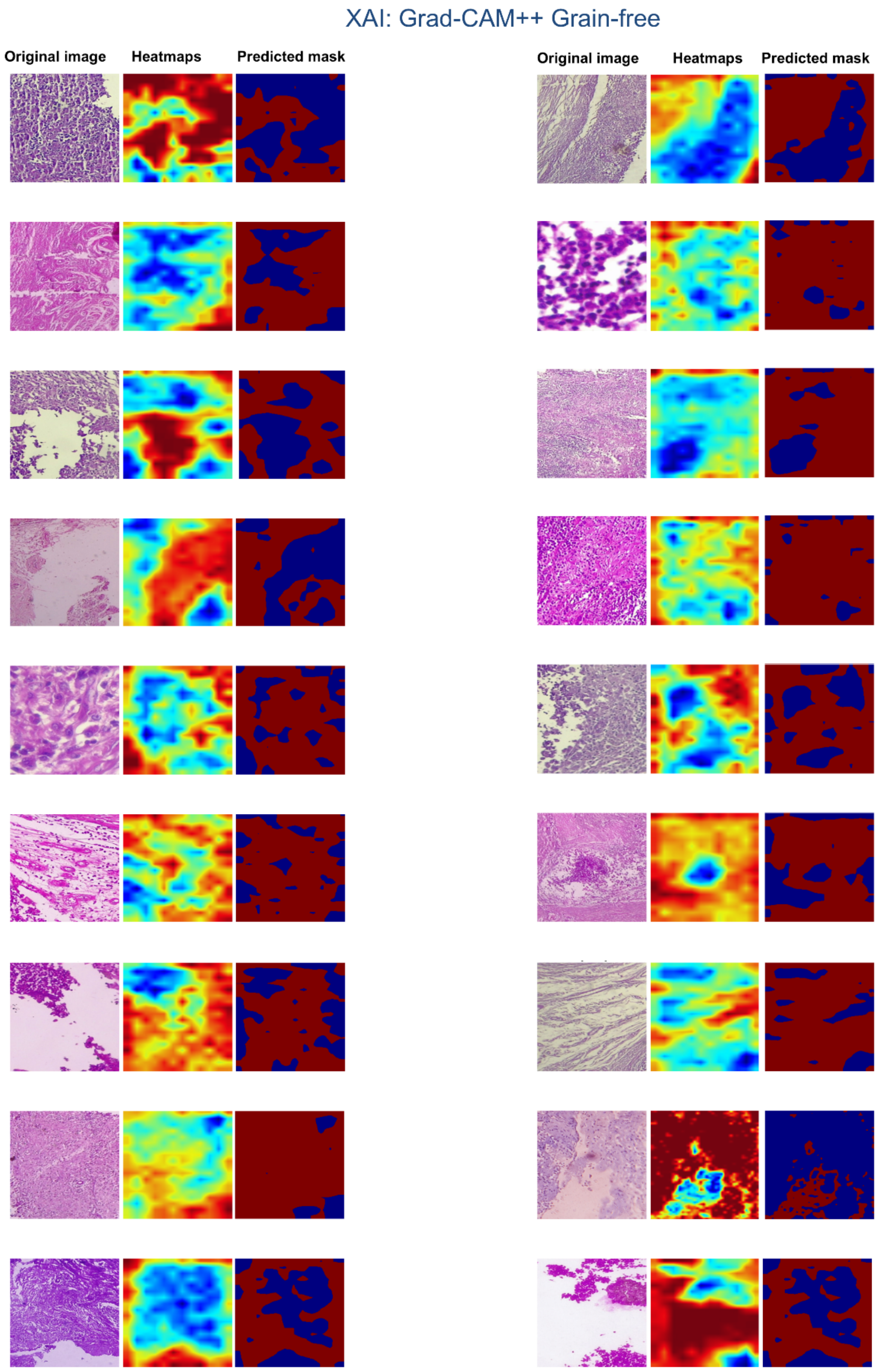


**Supplementary Figure S18: Grad-CAM++ Saliency Analysis for Explainable Grain-Free Image Classification.** This figure showcases representative examples of Grad-CAM++ saliency analysis applied to challenging grain-free images, serving as an Explainable AI task to demonstrate the visual interpretability of the classifier's decision-making. For each grain-free sample, the figure presents the original grain-free image, followed by its continuous-valued saliency heat mask (where warmer colors highlight regions most informative for classification), and finally, a binary saliency classification mask derived from the continuous heat map. This allows for critical insight into how the model makes decisions even when traditional diagnostic features (grains) are absent.
